# Commensal-derived succinate enhances tuft cell specification and suppresses ileal inflammation

**DOI:** 10.1101/776724

**Authors:** Amrita Banerjee, Charles A. Herring, Hyeyon Kim, Bob Chen, Alan J. Simmons, Austin N. Southard-Smith, James R. White, Marisol A. Ramirez Solano, Elizabeth A. Scoville, Qi Liu, M. Kay Washington, Ken S. Lau

## Abstract

Longitudinal analysis of Crohn’s disease (CD) incidence has identified an inverse correlation with helminth infestation and recent studies have revealed that intestinal tuft cell hyperplasia is critical for helminth response. Tuft cell frequency was decreased in the inflamed ilea of CD patients and a mouse model of TNFα-induced Crohn’s-like ileitis (TNFΔARE). Single-cell RNA sequencing paired with unbiased differential trajectory analysis of the tuft cell lineage in a genetic model of tuft cell hyperplasia (AtohKO) demonstrated that the tuft cell lineage had increased tricarboxylic acid (TCA) cycle gene signatures. Commensal microbiome-derived succinate was detected in the ileal lumen of these animals while microbiome depletion suppressed tuft cell hyperplasia. Therapeutic succinate treatment in TNFΔARE animals reduced pathology in correlation with induced tuft cell specification. We provide evidence implicating the modulatory role of intestinal tuft cells in chronic intestinal inflammation, which could facilitate leveraging this rare and elusive cell type for CD treatment.

## Introduction

Crohn’s disease (CD) is a relapsing-remitting Inflammatory Bowel Disease (IBD) characterized by chronic inflammation of the small intestine, with 60% of CD patients developing disease in the terminal ileum (Caprilli, 2008; Goulart *et al*, 2016). Despite the rising rate of IBD diagnoses, the etiology of CD remains unclear but is thought to be partially driven by a combination of a compromised barrier, a dysbiotic microbiome, and an altered immune response (Spalinger *et al*, 2014; Molodecky *et al*, 2012). Genome-wide association studies have clearly implicated epithelial-specific genes regulating microbial tolerance and clearance in IBD pathogenesis (Liu & Stappenbeck, 2016; de Lange *et al*, 2017; Liu *et al*, 2015; Barrett *et al*, 2009). In light of this, therapeutic strategies, including antibiotics and probiotics, to manipulate the microbiome for ameliorating disease activity have been attempted with limited success. Long-term remission is often not achieved with antibiotic therapy as patients often develop intolerance and other side effects to prolonged antibiotic regimens (Nitzan *et al*, 2016). Similarly, neither the VSL3 probiotic, which consists of eight lactic acid-producing bacterial species, nor fecal microbiome transplants have demonstrated the ability to induce response and remission in the majority of IBD patients, though no significant adverse effects have been reported from either treatment strategy (Bibiloni *et al*, 2005; Lopez & Grinspan, 2016). Since non-targeted microbiome therapies have had limited efficacy in achieving remission in CD, the development of targeted therapies may benefit CD treatment (Nitzan *et al*, 2016; Bibiloni *et al*, 2005).

Longitudinal analysis of global IBD incidence has identified an inverse correlation between the rates of communicable disease and autoimmune disorders (Molodecky *et al*, 2012; Saidel-Odes & Odes, 2014; Koloski *et al*, 2008). Known as the “hygiene hypothesis,” this phenomenon is thought to result from improved hygiene practices associated with decreased tolerance to environmental antigens (Saidel-Odes & Odes, 2014; Spalinger *et al*, 2014; de Silva MBBS & Korzenik, 2015). This paradoxical effect has led to emerging interest in the use of helminths, or parasitic worms, for the treatment of IBD (Helmby, 2015; Summers *et al*, 2003, 2005b, 2005a). Studies of the gut mucosa in major subsets of CD patients have demonstrated an increase in proinflammatory cytokines related to T-helper (Th)1 and Th17 cells, including interferon-γ and interleukin (IL)-17 as well as a commensurate decrease in Th2-associated cytokines, such as IL-25 and IL-13 (Su *et al*, 2013; Annunziato *et al*, 2015). An enhanced type 2 immune response, such as that seen in helminth infection, has been shown to suppress Th1 and Th17 activity (Su *et al*, 2013; Broadhurst *et al*, 2010). Clinical trial data from CD and ulcerative colitis (UC) patients have been inconclusive, as some trials have demonstrated decreased disease activity while others have been discontinued due to lack of efficacy (Summers *et al*, 2003, 2005b, 2005a). Moreover, helminth therapy has its drawbacks given prolonged infection can cause complications and, therefore, precision therapy using intermediary products may circumvent the majority of these issues (Summers *et al*, 2005a; Helmby, 2015).

In acute mouse models of infection, helminths colonize the proximal small intestine and provoke a type 2 immune response orchestrated by doublecortin-like kinase 1 (DCLK1)-positive epithelial tuft cells (Gerbe *et al*, 2016; von Moltke *et al*, 2016; Howitt *et al*, 2016). Through the release of IL-25, tuft cells promote their own specification via a positive feedback loop, and tuft cell hyperplasia is critical for pathogen clearance (Gerbe *et al*, 2016; von Moltke *et al*, 2016; Howitt *et al*, 2016). Tuft cell specification depends on genes, such as *Trpm5* and *Pou2f3*, which are canonical regulators of taste signal transduction (Gerbe *et al*, 2016; Howitt *et al*, 2016). Initially, studies of intestinal epithelial cell specification categorized DCLK1+ tuft cells within the secretory lineage, along with barrier-promoting goblet and Paneth cells, under the regulation of the master secretory transcription factor Atonal homolog 1 (*Atoh1*) (Gerbe *et al*, 2009, 2011). Recent studies have demonstrated that small intestinal tuft cells may have an alternative lineage specification route (Herring *et al*, 2018; Gracz *et al*, 2018) that may depend on cues from luminal microorganisms (Wilen *et al*, 2018). Despite the inverse relationship between incidence of parasitic infections and rates of IBD diagnoses, there is very little known about the functional role of tuft cells in human disease. In this study, we demonstrate that tuft cell specification is decreased in the inflamed small intestine of both human and mouse. By analyzing molecular pathway alterations in a lineage-specific manner during tuft cell hyperplasia, we identified commensal microbial-driven changes in metabolic pathways that promote tuft cell specification. Finally, metabolite-induced tuft cell hyperplasia in a mouse model of Crohn’s-like ileitis suppressed disease symptoms and restored epithelial architecture.

## Results

### Reduced tuft cell numbers are correlated to areas of ileal inflammation in human and mouse

Numerous studies have elucidated the critical role of tuft cells in driving a type 2 immune response against helminth infection (Helmby, 2015; Summers *et al*, 2005b, 2003; von Moltke *et al*, 2016; Howitt *et al*, 2016; Gerbe *et al*, 2016). Activation of the type 2 immune response has been implicated in the suppression of the proinflammatory environment in IBD patients (Helmby, 2015; Summers *et al*, 2005b, 2003). Thus, we first wanted to investigate the correlation between tuft cell numbers and local tissue inflammation in ileal specimens from CD patients, a study which to our knowledge had not been conducted. Tuft cell hyperplasia following acute helminth infection has been studied in the small intestine but not the colon (Howitt *et al*, 2016; von Moltke *et al*, 2016) and modes of tuft cell specification differ between the two tissues (Herring *et al*, 2018). Therefore, we restricted the sample inclusion criteria to inflamed ileal specimens (n = 14) from ileal CD patients compared to uninflamed normal ileal specimens (n = 11) collected from patients without a CD diagnosis, in order to limit our analysis to small intestinal tuft cells (Figure S1A).

One of the major reasons for the absence of studies on human tuft cells is the lack of validated markers. Previous attempts to use antibodies against DCLK1, a tuft cell-specific marker validated in the mouse intestine, to identify human tuft cells have not been successful (Gerbe *et al*, 2012). While others argue that current DCLK1 antibodies lack specificity against the human DCLK1 protein versus the mouse homolog, our single-cell RNA sequencing (scRNA-seq) data from the normal human ileum showed that the *DCLK1* gene was not significantly expressed in human tuft cells, demarcated by *TRPM5*, compared to murine tuft cells (Figure S1B). We have previously identified a double staining strategy of pEGFR(Y1068) and COX2 to specifically mark tuft cells in both mouse and human intestine (McKinley *et al*, 2017; Herring *et al*, 2018). Using this strategy, we observed double-positive pEGFR and COX2 cells in both the villi and crypts of Lieberkühn of the normal ileal epithelium (Figures S1C, 1A), that are distinct from single-positive pEGFR or COX2 cells in the lamina propria. Moreover, many of the double-positive cells possessed a prominent pEGFR-positive apical “tuft,” increasing the likelihood that these were genuine small intestinal tuft cells (Figure 1A).

**Figure 1.**
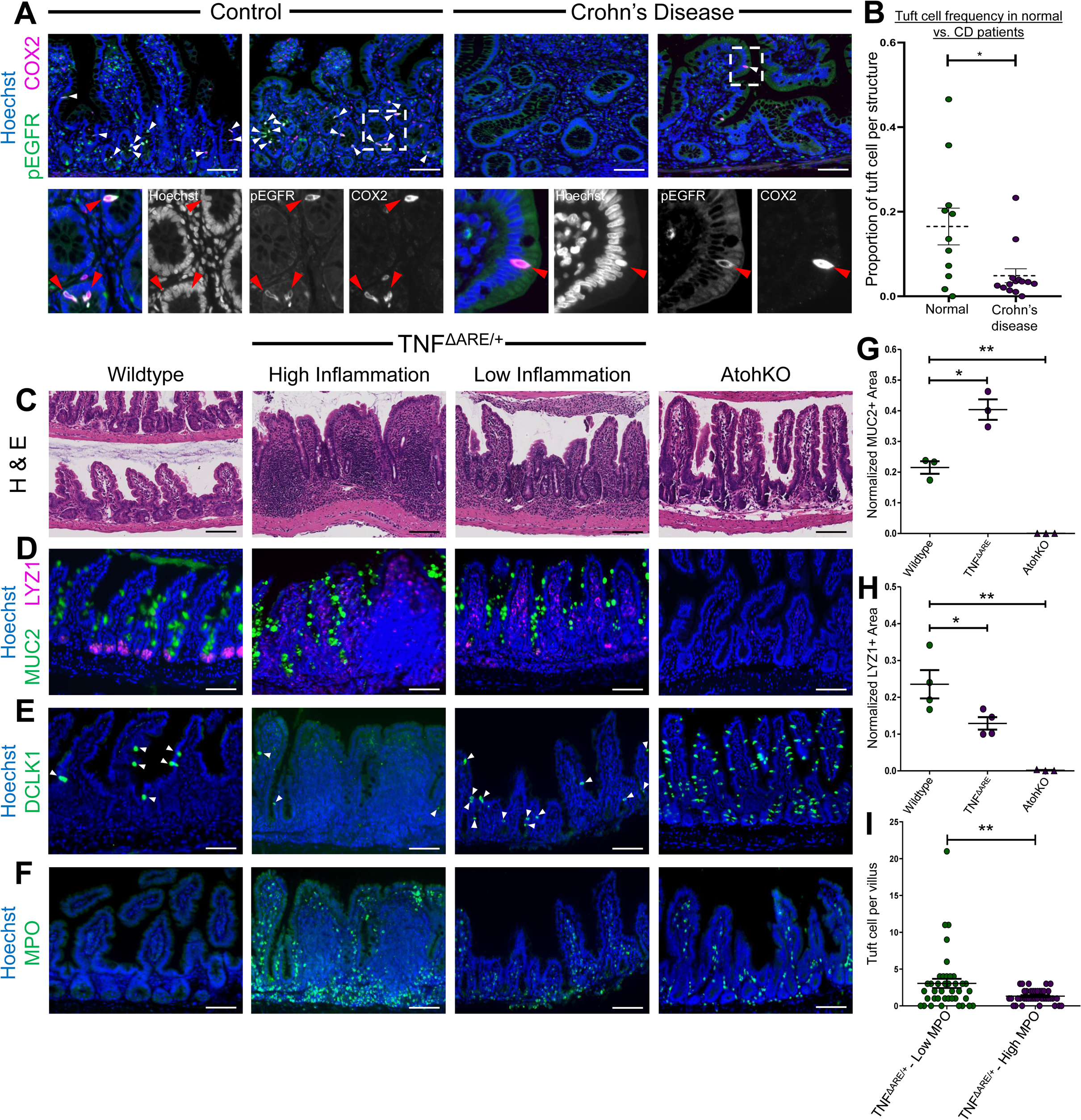
Human tuft cell number is decreased in biopsies of inflamed tissue from patients with ileal Crohn’s disease. **(A)** Immunofluorescence staining of pEGFR(Y1068) (green) and COX2 (magenta) in human ileal tissues from healthy and Crohn’s disease patients. Co-localization of pEGFR and COX2 is used to identify small intestinal tuft cells, demarcated by white arrows. Magnified inset of the epithelium shows that both markers are expressed in individual tuft cells, demarcated by red arrows. Representative images are shown from two separate normal or Crohn’s disease patients, respectively. Hoechst (blue) denotes nuclei, scale bar = 100 µm. **(B)** Quantification of pEGFR and COX2 double-positive tuft cells in normal (green) and Crohn’s disease (magenta) ileal samples. Each dot represents a separate sample. Error bars represent SEM for n = 11 normal and n = 14 Crohn’s disease samples, respectively. *p-value < 0.05 by t-test. **(C)** Histology of distal ileum from wildtype, low and high inflammation TNF^ΔARE/+^, and AtohKO animals. Scale bar = 100 µm. **(D)** Immunofluorescence imaging of MUC2 (green) and LYZ1 (magenta). Hoechst (blue) denotes nuclei, scale bar = 100 µm. **(E)** Immunofluorescence imaging of DCLK1 (green) and Hoechst (blue). Scale bar = 100 µm. **(F)** Immunofluorescence imaging of MPO (green) and Hoechst (blue). Scale bar = 100 µm. **(G)** Quantification of MUC2 staining normalized by Hoechst area in villi. Error bars represent SEM from n = 3 mice per condition. **p-value < 0.01, *p-value < 0.05 by t-test. **(H)** Quantification of LYZ1 staining normalized by Hoechst area per crypt. Error bars represent SEM from n = 4 wildtype and TNF^ΔARE/+^ mice and n = 3 AtohKO animals. **p-value < 0.01, *p-value < 0.05. **(I)** Tuft cell number per villi in the TNF^ΔARE/+^ ileum stratified by MPO+ neutrophils. Low MPO < 35 neutrophils and High MPO ≥ 35 neutrophils. Error bars represent SEM for n = 40 villi per condition across 6 TNF^ΔARE/+^ animals. **p-value < 0.01 by t-test.

We applied this strategy to detect tuft cells from inflamed ileal tissues of CD patients. As expected, inflammation in ileal CD samples was heterogenous and characterized by severe villus blunting (Figure S1D). Consistent with previous reports, MUC2+ goblet cells were increased in the inflamed epithelium, while LYZ+ Paneth cells were decreased (Figure S1E) (VanDussen *et al*, 2014; Antoni *et al*, 2014; Wehkamp *et al*, 2007, 2016; Erben *et al*, 2016). The few Paneth cells remaining in inflamed regions contained more diffuse apical, lysozyme-positive granules, consistent with known Paneth cell phenotypes in CD (VanDussen *et al*, 2014). LYZ expression was increased in the lamina propria in inflamed tissue, most likely from active immune cells (Figure S1E). Quantification of tuft cells, detected by co-staining of pEGFR and COX2, revealed a significant reduction in ileal specimens from CD patients (Figures 1A-B). Within our pool of CD patients, there was no further, progressive decrease in tuft cell number with increased disease severity (Figure S1F). In certain CD specimens, particularly in regions with less disease involvement and more organized tissue architecture, tuft cells could still be detected by pEGFR and COX2 staining (Figure 1A). From these results, we speculate that suppression of tuft cell specification may contribute to the loss of inflammation control in CD and thus may be associated with disease development and/or progression.

To assess tuft cell specification in a more controlled manner, we used the TNF^ΔARE/+^ mouse model, which, by the deletion of an AU rich element (ΔARE) in the gene encoding the proinflammatory cytokine tumor necrosis factor alpha (*Tnfa*), has increased levels of the TNF-α mRNA and develops Crohn’s-like ileitis by two to three months of age (Kontoyiannis *et al*, 2002, 1999; Erben *et al*, 2016). The TNF^ΔARE/+^ model mimics many features of human ileal CD, including dependence on TNF-α and microbiome dysbiosis (Roulis et al, 2016b; Goulart et al, 2016). We observed histological changes in the terminal ileum of four-month-old TNF^ΔARE/+^ animals, characterized by distorted crypt structure and blunted villi compared to wildtype littermates (Figure 1C). The number of LYZ1+ Paneth cells was decreased while the remaining ones exhibited diffuse LYZ1 staining, suggesting impaired Paneth cell function (Figures 1D, 1H, S2B) (VanDussen et al, 2014; Wehkamp et al, 2005). The numbers of LYZ1+ cells in the lamina propria and MUC2+ goblet cells in the epithelium were increased (Figures 1D, 1G, S2A), which bear resemblance to the ilea of human CD patients (Wehkamp et al, 2005; Erben et al, 2016). Increased immune cell infiltration can be observed by myeloperoxidase (MPO)-positive neutrophils in the lamina propria of inflamed tissues compared with uninflamed controls (Figure 1F).

Similar to the heterogeneity observed in specimens from CD patients, we were able to identify both highly inflamed and less inflamed regions within the ilea of TNF^ΔARE/+^ mice (Figure S2C). We performed spatially-resolved analysis to determine the relationship between tuft cell numbers and inflammation by quantifying the number of DCLK1+ tuft cells in the epithelium and infiltrating MPO+ neutrophils in the lamina propria on a per-villus basis (Figure S2D). Consistent with human intestinal phenotypes, regions classified as highly inflamed with >35 MPO+ cells/villus were characterized by severe villus blunting and distortion, while less inflamed regions (0-35 MPO+ cells/villus) possessed normal crypt-villus architecture and resembled healthy wildtype controls (<10 MPO+ cells/villus) (Figures S2C-S2D). Within the same TNF^ΔARE/+^ animal, tuft cell numbers were significantly decreased in highly inflamed regions compared to less inflamed regions (Figures 1E, 1I). The number of tuft cells in less inflamed regions was increased beyond the normal number of tuft cells found in the healthy ileum (Figure 1E). These results are consistent with our observations in human ileal biopsies and suggests that increasing tuft cell specification may suppress ileal inflammation.

### Tuft cells are specified outside of the secretory lineage in ileal inflammation

Based on our observations in human and mouse inflammation, we hypothesized that increasing tuft cell specification may potentially mitigate CD symptoms. Therefore, we sought to better understand the signals and pathways regulating tuft cell specification, specifically in the context of perturbed microbiome-epithelium interaction in IBD (Boyapati et al, 2015; Roulis et al, 2016a). However, the regional heterogeneity of inflammation and tuft cell specification in the TNF^ΔARE/+^ model precludes most systematic analysis. To uncover the signals governing tuft cell specification in a more tractable manner, we generated a genetically inducible model of tuft cell hyperplasia that is homogeneous throughout the entire ileum. The *Lrig1^CreERT2^* driver was crossed to *Atoh1^loxP/loxP^* mice to generate *Lrig1^CreERT2/+^; Atoh1^fl/fl^* (AtohKO) animals, where the administration of tamoxifen drove recombination of *Atoh1* in *Lrig1*-expressing stem cells (Herring *et al*, 2018). While AtohKO animals exhibited decreased weight gain (SF 2E), they possessed relatively normal crypt-villus architecture, as seen by H&E staining (Figure 1C). Quantification of immunofluorescence staining confirmed that AtohKO animals lacked MUC2+ goblet cells and LYZ1+ Paneth cells (Figures 1D, 1G-1H), consistent with their classification in the *Atoh1*-dependent secretory lineage (Vandussen & Samuelson, 2010; Shroyer *et al*, 2007; Noah *et al*, 2011). However, in contrast to the established hierarchy, we observed a significant increase in DCLK1+ tuft cells throughout the small intestine (Figure 1E), which supports more recent studies suggesting tuft cells can be specified outside the *Atoh1*-dependent secretory lineage (Herring *et al*, 2018; Gracz *et al*, 2018). In addition, we observed an increased presence of MPO+ neutrophils in the villi of AtohKO animals, reflecting a baseline level of inflammation, similar to the less inflamed regions of TNF^ΔARE/+^ animals, potentially due to goblet and Paneth cell loss (Figure 1F). The uniformity of these phenotypes throughout the ilea of AtohKO mice allows us to investigate mechanisms driving tuft cell specification in an *in vivo* model with a physiologically intact microbiome and immune system.

In order to decipher lineage-specific alterations that drive tuft cell specification, we generated scRNA-seq data from wildtype, TNF^ΔARE^*^/+^*, and AtohKO ilea using the inDrop platform (Klein *et al*, 2015), by enriching for epithelial crypts and generating single-cell suspensions via a cold protease dissociation protocol (Adam *et al*, 2017). Following data processing and quality control, we obtained a 6,932-cell wildtype scRNA-seq dataset from six biological replicates (Figures S3A, S3D, S3G) and performed t-SNE analysis on 1,450 randomly selected cells from the entire complement of datapoints (Figure 2A). Clustering analysis demonstrated that the wildtype scRNA-seq dataset contained the correct proportion of expected cell types, including stem and progenitor cells, enterocytes, goblet cells, Paneth cells, enteroendocrine cells, and tuft cells (Figures 2A, S3D, S4D, S4A). To compare across conditions, 1,450 datapoints were randomly selected for further analysis from the 3,401-cell TNF^ΔARE/+^ (Figures 2B, S3B, S3H) and 2,456-cell AtohKO scRNA-seq datasets (Figures 2C, S3C, S3I). In the TNF^ΔARE/+^ dataset, while all the cell types were represented, the Paneth cell population was greatly diminished, consistent with results observed by immunofluorescence imaging (Figures 2B, S3E, S4B, S4E). Quantification of *Dclk1*-expressing cells indicated that the tuft cell population is not significantly different in the TNF^ΔARE^*^/+^* dataset (Figures 2D, S4E), which may be accounted for by the heterogeneity observed in the inflamed ileum. Given the loss of spatial resolution subsequent to single-cell dissociation, it is reasonable that a global approach such as scRNA-seq cannot accurately capture the spatial heterogeneity of the TNF^ΔARE^*^/+^* model. The AtohKO scRNA-seq data confirmed the loss of goblet, Paneth, and enteroendocrine lineages following *Atoh1* recombination, consistent with their *Atoh1*-dependent secretory origins (Figures 2C, S3F, S4C, S4F). Compared to the wildtype tuft cell cluster, the AtohKO tuft cell population was significantly expanded (Figures 2C-2D).

**Figure 2.**
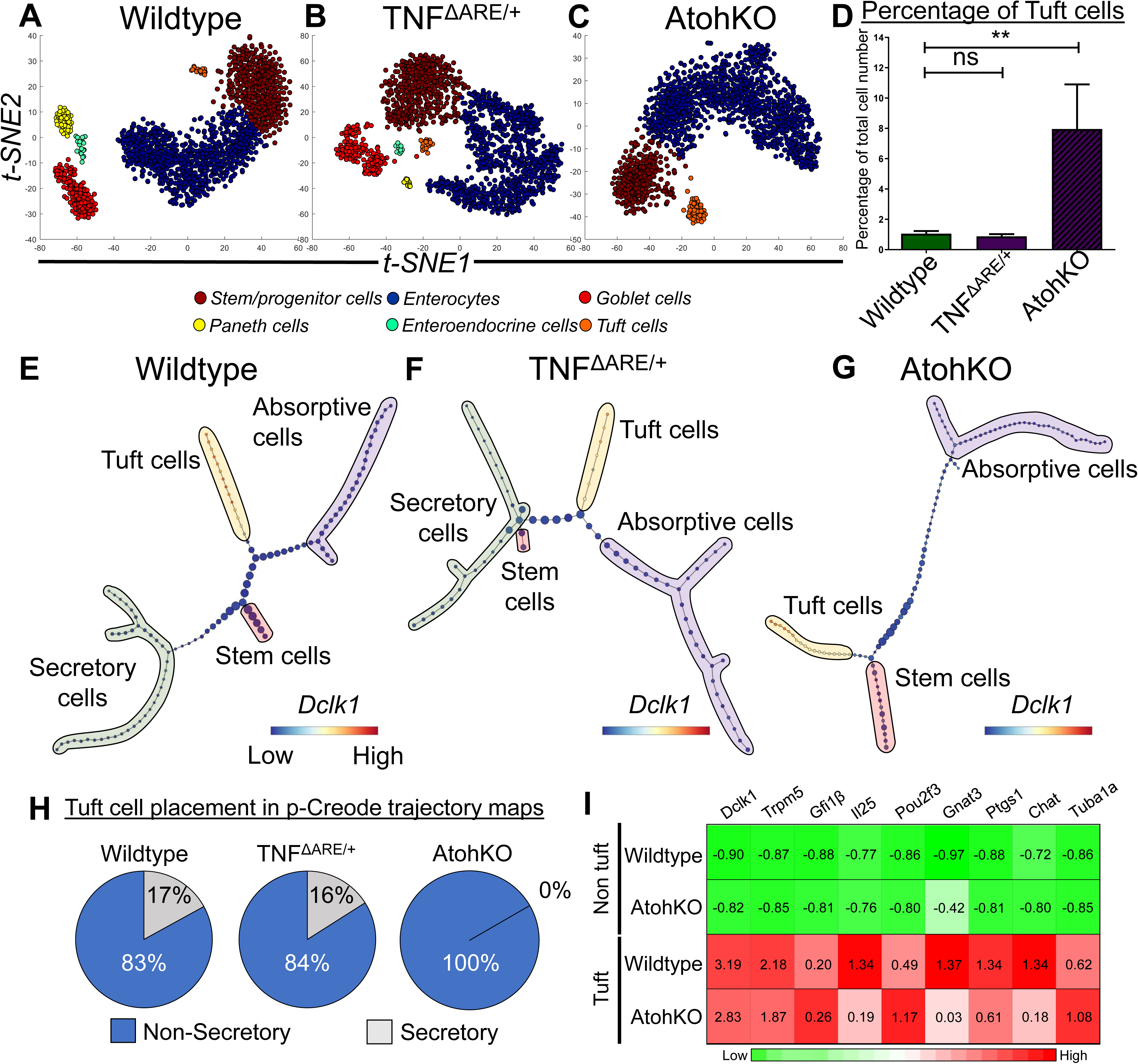
p-Creode trajectory analysis of ileal epithelial scRNA-seq data supports an alternate origin for small intestinal tuft cells. **(A-C)** t-SNE analysis of scRNA-seq data generated from **(A)** wildtype, **(B)** TNF^ΔARE/+^, and **(C)** AtohKO ileal epithelium. Cell type clusters, including goblet cells (red), Paneth cells (yellow), enteroendocrine cells (green), tuft cells (orange), enterocytes (dark blue), and stem/progenitor cells (brown), were identified by k-means clustering and manually annotated. Each t-SNE plot depicts 1,450 randomly selected datapoints from their corresponding complete dataset and each datapoint represents a single cell. **(D)** Quantification of tuft cell percentage within the scRNA-seq datasets of wildtype, TNF^ΔARE/+^, and AtohKO ileal epithelium. Error bars are generated from n = 6 wildtype replicates, n = 3 TNF^ΔARE/+^ replicates, and n =3 AtohKO replicates. **p-value < 0.01, *p-value < 0.05, and ns (not significant) by t-test. **(E-G)** p-Creode analysis of scRNA-seq datasets shown in **A-C**, depicting the most representative topology map over n = 100 runs for **(E)** wildtype, **(F)** TNF^ΔARE/+^, and **(G)** AtohKO datasets. Graph overlay depicts ArcSinh-scaled *Dclk1* gene expression data. Cell lineages, including secretory cells (green), absorptive cells (purple), tuft cells (orange), and stem cells (red), were manually labelled. Node size represents cell state density and each edge represents cell state transitions. **(H)** Quantification of n = 100 p-Creode maps for wildtype, TNF^ΔARE/+^, and AtohKO datasets, respectively. Tuft cell placement was classified as secretory (grey) when the tuft cell lineage shared a trajectory with the secretory lineage and as non-secretory (blue) when the tuft cells and absorptive lineage shared a trajectory. **(I)** Heatmap depicting z-score normalized expression of tuft cell gene signature in tuft and non-tuft cell populations in both wildtype and AtohKO datasets. All values are statistically significant, ***p-value < 0.001 by t-test.

We analyzed tuft cell specification pathways using the p-Creode algorithm to produce trajectory representations of our scRNA-seq datasets (Herring *et al*, 2018). The wildtype p-Creode map originated from the stem cell lineage and bifurcated into the secretory and absorptive lineages (Figures 2E, S5A). As expected, goblet and Paneth cells originated from a common secretory progenitor before diverging into two distinct lineages (Figure S5A). In contrast, the tuft cell lineage shared a specification trajectory with absorptive cells, rather than the secretory cells (Figures 2E, S5A). In order to evaluate the robustness of this map, we generated 100 p-Creode graphs by randomly sampling the wildtype scRNA-seq dataset and quantified tuft cell placement. p-Creode maps were classified as “secretory” when the tuft cells were grouped with lineages containing goblet cells and as “non-secretory” otherwise. Tuft cell placement was non-secretory in 83% of wildtype trajectories and secretory in the remaining 17% (Figure 2H). These results supported recent findings regarding an alternative origin of small intestinal tuft cells that is separate from the *Atoh1*-dependent secretory lineages (Herring *et al*, 2018; Gracz *et al*, 2018). p-Creode analysis of the TNF^ΔARE^*^/+^* scRNA-seq dataset illustrated that, even under inflammatory conditions, tuft cells shared a trajectory with the absorptive lineage (Figures 2F, S5B). Quantification of 100 p-Creode runs showed that tuft cell placement was non-secretory in 84% of TNF^ΔARE/+^ maps and secretory in 16% of maps (Figure 2H). Finally, due to the loss of the secretory lineages in the AtohKO model, all 100 p-Creode graphs generated from the corresponding scRNA-seq data depicted tuft cells and absorptive cells originating from a common progenitor (Figures 2G-2H, S5C). Gene expression of tuft cell regulators, including *Pou2f3*, *Ptgs1*, *Ptgs2*, *Sox4*, *Sox9*, and *Trpm5*, was confirmed to be expressed in the tuft cell lineage in wildtype, TNF^ΔARE/+^, and AtohKO p-Creode topologies (Figure S6A-S6C). Moreover we confirmed that tuft cells in the AtohKO small intestine express the tuft cell gene signature (von Moltke *et al*, 2016) while non-tuft cells do not, similar to the wildtype condition (Figures 2I, S4G).

We repeated the p-Creode analysis to include rare enteroendocrine cells in the wildtype p-Creode topology and observed that, while enteroendocrine cells segregate with secretory cells (Figures S7A, S7C), tuft cells still by-and-large share a trajectory with the absorptive cells (Figures S7B). To confirm these results with an alternative dataset, we re-analyzed a 7,000+-cell scRNA-seq dataset generated using 10X Genomics by Aviv Regev’s group (GSE92332), from which we were able to reproduce the expected distribution of cell types (Figure S8A-S8C) (Haber *et al*, 2017). p-Creode analysis demonstrated cell differentiation originated from stem cells, and bifurcated into the absorptive and secretory cells, which further diverged into the Paneth and goblet cell lineages (Figures S8D, S8F). In addition, we observed non-secretory placement of the tuft cell lineage with the absorptive lineage in 68% of p-Creode topologies and 32% placement with the secretory lineages (Figures S8E). Slight discrepancies in the results of the two analyses may be accounted for by technical differences between the datasets in regards to cell isolation, library preparation, or data processing procedures. Nevertheless, tuft cell specification from a non-secretory lineage was a robust and consistent feature of the wildtype small intestine across multiple datasets. Moreover, given that the AtohKO tuft cell population expressed a similar signature as wildtype tuft cells, this model can reliably be used to study pathways which govern tuft cell specification.

### Alterations in TCA metabolic pathways along the tuft cell trajectory are associated with induced tuft cell specification

p-Creode trajectory mapping enables analysis of gene expression changes in a lineage-specific manner (Liu *et al*, 2018; Herring *et al*, 2018). Within the intestinal epithelium, expression of *Dclk1* gradually increased in the tuft cell lineage but was undetectable in the enterocyte and goblet cell lineage (Figure S9A). Conversely, *Muc2* and *Krt20* expression gradually increased as goblet cells and enterocytes differentiate in their respective lineages (Figure S9A). To identify pathways that induce tuft cell differentiation, we focused our analysis on dynamic alterations in gene expression along the tuft cell lineage between the wildtype and AtohKO intestinal epithelium. This approach circumvented technical batch effects, since the dynamics of gene expression along a trajectory is self-contained within individual analyses.

We aimed to identify genes that switch their expression dynamics between the wildtype and AtohKO conditions. First, different types of gene dynamics in the tuft cell lineage were broadly classified into four categories, as well as a fifth category of unchanged or “flat” dynamics (Figure 3A). Group one genes, such as *Soux*, trended upward along pseudotime of the stem-to-tuft cell trajectory, while group four genes, including *Rps6*, trended downwards (Figure S9B). Group one genes included known tuft cell marker genes that are upregulated during differentiation (for instance, *Ptgs1* and *Sox9*) (McKinley *et al*, 2017), while group four genes included stem cell markers that are downregulated during differentiation. Intermediate genes that trended upwards but returned to a lower baseline or those that trended downwards but returned to a higher baseline were categorized into groups two and three, respectively (Figure S9B). When all expressed genes between the wildtype and AtohKO were visualized within these categories, a broad expansion of group two genes was observed upon loss of *Atoh1*, implicating changes in lineage-specific gene expression dynamics (Figure 3A). To identify pathways that induced tuft cell specification in an unbiased manner, we extracted 1,755 genes that were positively enriched in the AtohKO group, namely, those that switched categories from a lower category in the wildtype data to a higher category in AtohKO (Figure S9C). Over-representation analysis of positively enriched genes in the AtohKO epithelium identified pathways related to the tricarboxylic acid (TCA) cycle and oxidative phosphorylation based on KEGG (Figure 3B), Wiki pathways (Figure S9D), and Reactome analyses (Figures S9E). We repeated this analysis by grouping the dynamic trends into either “up” or “down” and again identified genes that were positively enriched in the AtohKO trajectory. This coarse grain analysis produced similar results as before, as over-representation analysis identified enrichment for metabolic-associated processes, including the TCA cycle and the electron transport chain (Figures S9F-S9H).

**Figure 3.**
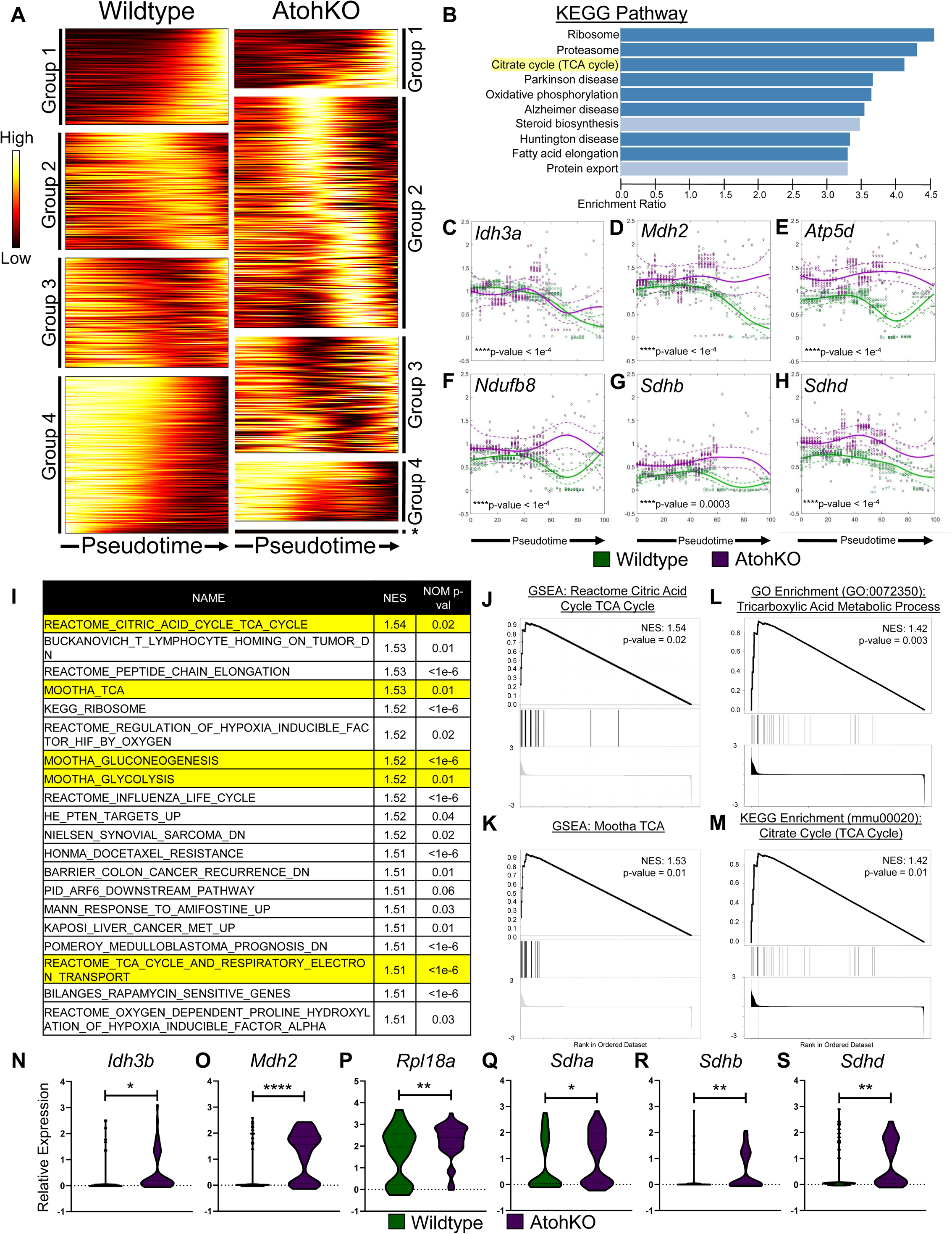
Analysis of AtohKO tuft cell gene expression identified upregulation in metabolic pathways. **(A)** Heatmap of gene expression trends across pseudotime in tuft cell lineage from wildtype and AtohKO p-Creode topologies. Genes were clustered by their dynamics – Groups 1-2 included upregulated genes and Groups 3-4 included downregulated genes. Group 5 (**\***) in AtohKO consisted of genes that were 0 expression. **(B)** KEGG enrichment bar plots for genes that class switch from lower order in wildtype tuft cells to higher order in AtohKO tuft cells. Functional groups were ordered by NES. **(C-H)** Trend dynamics along pseudotime of citrate cycle-related genes for the wildtype (green) and AtohKO (magenta) tuft cell lineages. Solid lines represents wildtype (green) and AtohKO (magenta) gene expression trends from 10 representative p-Creode graphs. Raw data is shown for wildtype (green circles) and AtohKO (magenta diamonds). Confidence interval of raw data was depicted by dashed lines for wildtype (green) and AtohKO (magenta). Dynamic time warping was used to fit the wildtype and AtohKO tuft cell data to the same scale. Statistical analysis of trend differences and consensus alignment was performed between conditions, ****p < 0.0001 by t-test. **(I)** Gene set enrichment analysis of median difference between wildtype (n = 58 cells) and AtohKO (n = 64) tuft cell populations. Top 20 gene sets from positive gene enrichment are ranked by the normalized enrichment score (NES) and p-value. Yellow highlighted gene sets are related to the citric acid cycle and metabolism pathways. **(J-K)** Positive enrichment plots for the gene sets with the highest NES based on GSEA. **(L-M)** Pathway analysis between wildtype and AtohKO tuft cells for **(L)** gene ontology and **(M)** KEGG analysis shows positive enrichment for the Tricarboxylic or citrate acid cycle Metabolic Process. **(N-S)** Relative expression of TCA cycle genes in wildtype (green) and AtohKO (magenta) tuft cells. Error bars represent SEM from wildtype (n = 58 cells) and AtohKO (n = 64 cells). ****p < 0.0001, < 0.001, **p < 0.01, and *p < 0.05 by t-test.

The TCA cycle converts glycolysis products into NADH+, which is then shuttled to the electron transport chain for ATP production (Mills & O’neill, 2014). Alterations in the wildtype (green) and AtohKO (magenta) tuft cell lineages of selected pathway genes based on KEGG analysis were directly visualized by plotting expression trends fit to raw data from ten representative p-Creode maps (Figure 3C-3H). As benchmarks, we observed that dynamics of tuft and stem cell signature genes were unaltered between wildtype and AtohKO trajectories (Figures S9I). Our visualization method confirmed that regulators of the tuft cell lineage, including *Dclk1* and *Trpm5*, trended upward, while stem or progenitor cell genes, *Myc* and *Pcna*, trended downward during the differentiation trajectory (Figures S9I). In contrast, the TCA cycle enzyme malate dehydrogenase (*Mdh2*) trended down along the wildtype tuft cell differentiation trajectory, while its expression remained constant in the AtohKO tuft cell trajectory (Figure 3D). Similarly, other TCA enzymes, such as *Idh3a* (Figure 3C)*, Sdhb* (Figure 3G), and *Sdhd* (Figure 3H), all switched to more positive dynamic trends along the AtohKO tuft cell trajectory compared to the wildtype tuft cell trajectory. Other related genes, including those coding NADH dehydrogenases and ATP synthases also switched to more positive dynamic trends in the AtohKO tuft cell lineage (Figures 3E-3F, S11). Analysis of altered dynamics suggests that tuft cell hyperplasia in the AtohKO model is accompanied by increased TCA cycle and downstream metabolic activities along the tuft cell specification trajectory.

As a confirmatory method, we grouped cells within the tuft cell trajectory and performed standard differential expression analysis between wildtype and AtohKO cell populations. Standard gene set enrichment analysis (GSEA) (Subramanian *et al*, 2005) performed on genes upregulated in AtohKO tuft cells identified positive enrichment for TCA cycle genes (Figures 3I, S11A). Enrichment plots for the two GSEA gene sets with the highest normalized enrichment score (NES), “Reactome_Citric_Acid_Cycle_TCA_Cycle” and “Mootha_TCA” are shown in Figure 3 and the expression of highly enriched genes (Figure S11F) from the former gene set was compared between wildtype and AtohKO tuft cells (Figures 3J-3K). TCA cycle-related enzymes *Idh3b*, *Mdh2*, *Sdha*, *Sdhb,* and *Sdhd,* as well as the ribosomal protein gene *Rpl18a,* were all significantly higher in AtohKO (magenta) tuft cells compared to wildtype tuft cells (green) (Figure 3N-3S). Expression of other TCA cycle enzymes, including *Citrate synthase (Cs), Idh3a*, *Idh3g*, *Ogdh*, and *Sdhc*, while not statistically significant, also trended upward in AtohKO tuft cells (Figures S11H-S11L). Additional GSEA over Gene ontology (Figures 3L, S11D), KEGG (Figures 3M, S11B), PANTHER (Figure S11C), and Wiki pathways (Figure S11E) gene sets also identified that TCA cycle-associated genes were upregulated in the AtohKO tuft cells, further suggesting activation of metabolic pathways in this cell population.

### Non-parasite-derived sources of succinate drive tuft cell specification

While metabolic pathways were upregulated in the AtohKO tuft cell lineage, it was unclear whether these changes arose from epithelial cell-intrinsic function of the ATOH1 transcription factor or other non-cell autonomous mechanisms. The AtohKO epithelium lacked barrier-regulating goblet and Paneth cells, which may have led to changes in the microbiome and an influx of tuft cell-promoting commensal-derived metabolites. To test the necessity of the microbiome in driving tuft cell hyperplasia, we induced *Atoh1* recombination in enteroid cultures in a sterile environment (Sato *et al*, 2011). Vehicle-treated enteroids had robust expression of LYZ1+ Paneth cells, MUC2+ goblet cells, and DCLK1+ tuft cells, as expected (Figures S12A-S12B). Similar to the *in vivo* condition, loss of *Atoh1* via Cre recombination led to an absence of Paneth and goblet cells (Figures S12C-S12D). However, tuft cell hyperplasia was not observed in sterile enteroids (Figures S12C-S12D), which implies tuft cell hyperplasia in the *in vivo* AtohKO model may arise via a microbiome-dependent mechanism.

To assess the necessity of the microbiome for inducing tuft cell hyperplasia *in vivo*, we used an antibiotic cocktail, consisting of kanamycin, metronidazole, gentamicin, colistin sulfate, and vancomycin, to deplete a broad range of gram-positive and gram-negative bacteria with minimal impact on overall health (Meng *et al*, 2007; Julia *et al*, 2000) (Figures S12E). Loss of *Atoh1* in this context resulted in depletion of LYZ1+ Paneth cells (Figures 4AI-4AIV), but antibiotic administration significantly suppressed the resulting tuft cell hyperplasia (Figures 4BII-4BIV) in comparison to the untreated control (Figure 4B I). Similar results were observed in AtohKO animals treated with ampicillin, which targets only gram-negative bacteria (Figure S12F). The effects of antibiotic-based microbiome depletion were dose-dependent, with a general suppression of tuft cell hyperplasia correlated to increasing antibiotic concentration (Figures 4BII - 4BIV). To identify the pathways associated with this suppression, we generated scRNA-seq datasets from AtohKO animals on a low to mid dose antibiotic regimen, where tuft cells were still being specified, albeit at a lower frequency (Figures 4BII-4BIII). Based on genes gleaned from prior GSEA analysis, we compared the expression of TCA cycle-related genes in the AtohKO tuft cells from vehicle and antibiotic-treated mice. TCA cycle enzymes *Idh3b, Mdh2*, *Shda, Sdhb,* and *Sdhd*, which were upregulated in AtohKO tuft cells, were significantly decreased following antibiotic treatment (Figures 4C-4G). Additionally, TCA cycle-associated gene *Rpl18a*, which was highly enriched in the AtohKO tuft cells, was decreased in the antibiotic-treated datasets (Figure 4H). *Idh3a, Idh3g,* and *Sdhc*, which trended upward in the AtohKO tuft cells, trended downward in the antibiotic-treated AtohKO condition (Figures S12H-S12I, S12K), while *Cs* and *Ogdh* were increased (Figures S12G, S12J). These results indicated that extrinsic signals from the intestinal microbiome drive gene expression changes in the AtohKO tuft cell lineage to induce tuft cell hyperplasia *in vivo*.

**Figure 4.**
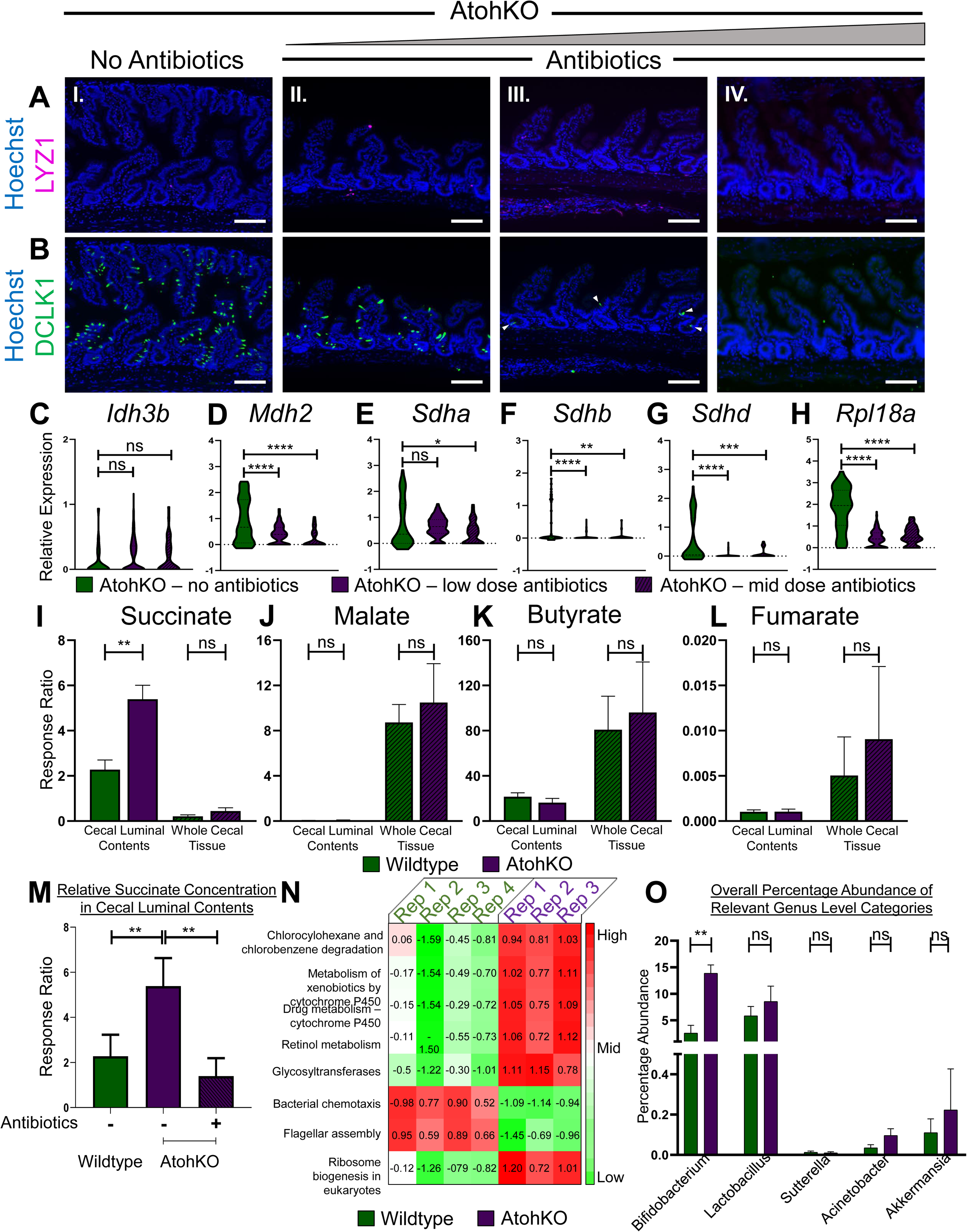
*In vivo* tuft cell hyperplasia in the AtohKO small intestine is microbiome-dependent. **(A-B)** Representative immunofluorescence staining of **(A)** LYZ1 (magenta) and **(B)** DCLK1 (green) in the AtohKO ileum, **(I)** without antibiotics and with **(II)** low dose, **(III)** mid dose, and **(IV)** high dose antibiotics. Hoechst (blue) denotes nuclei, scale bar = 100 µm. **(C-H)** Relative expression of TCA cycle genes in AtohKO – no antibiotics (green), AtohKO – low dose antibiotics (magenta), and AtohKO – mid dose antibiotics (magenta, dashed) tuft cells. Error bars represent SEM from untreated AtohKO (n = 25 cells), low-dose antibiotic-treated AtohKO (n = 202 cells), and mid-dose antibiotic-treated AtohKO (n = 36 cells). ***p-value < 0.001, *p-value < 0.05, and not significant (ns) by t-test. **(I-H)** Relative concentration of short chain fatty acids, **(I)** succinate, **(J)** malate, **(K)** butyrate, and **(L)** fumarate in cecal luminal contents and whole tissue from wildtype (green) and AtohKO (magenta) animals. Error bars represent SEM across n = 5 wildtype and n = 3 AtohKO replicates. **p-value < 0.01 and not significant (ns) by t-test. **(M)** Relative succinate concentration in cecal luminal contents among wildtype (green), untreated AtohKO (magenta), and antibiotic-treated AtohKO (magenta, dashed) animals. Error bars represent SEM across n = 5 wildtype, n = 3 AtohKO, and n = 3 antibiotic-treated AtohKO replicates. **p-value < 0.01 by t-test. **(N)** PICRUSt functional analysis of 16s profiles from wildtype (green) and AtohKO (magenta) replicates. Heatmap of z-score normalized relative abundances of statistically significant predicted functional categories from wildtype and AtohKO 16s data. *p-value < 0.05 by test. **(O)** Relative abundance of relevant genus level categories contributing to the “Chlorocyclohexane and chlorobenzene degradation category” between wildtype and AtohKO microbiome profiles. Error bars represent SEM from n = 4 wildtype and n = 3 AtohKO replicates. **p-value < 0.01 and not significant (ns) by t-test.

In order to investigate the microbial-derived signals driving the increased tuft cell specification, we sought to characterize the differences in (1) commensal-derived metabolites and (2) relative microbial abundance in the lumen of wildtype and AtohKO animals. We used O-benzylhydroxylamine (O-BHA) derivatization to analyze metabolite levels in cecal luminal contents and cecal tissue (Tan *et al*, 2014). Short chain fatty acids, such as malate and butyrate were not significantly increased in the AtohKO condition (Figures 4J, 4K). However, O-BHA analysis revealed that the relative concentration of succinate was significantly increased in the AtohKO cecal luminal contents but not in cecal tissue, compared to the wildtype condition (Figure 4I). Succinate or succinic acid is a metabolic intermediate in the TCA cycle and is converted into fumarate by the enzyme succinate dehydrogenase (Mills & O’neill, 2014; Ryan *et al*, 2019). Fumarate levels were negligible in both the wildtype and AtohKO lumen but increased in the AtohKO tissue, suggesting metabolic processing of luminal succinate (Figure 4L). Moreover, the disparity in succinate concentration between luminal contents and whole tissue were indicative of a commensal microbiome, rather than a host-derived, origin for succinate (Figure 4I). To support this hypothesis, we repeated the analysis in the AtohKO condition following microbiome depletion. Succinate levels were significantly decreased in the antibiotic-treated AtohKO cecal luminal contents, confirming that the commensal microbiome was primarily responsible for succinate production in this model (Figure 4M).

Previously published work has demonstrated that parasite-derived succinate, downstream of helminth infection, can drive tuft cell hyperplasia and induce the type 2 immune response necessary for worm extrusion (Nadjsombati *et al*, 2018; Lei *et al*, 2018; Schneider *et al*, 2018). We confirmed that succinate administration induces tuft cell hyperplasia (Figure S12L), while major basic protein-positive eosinophils (Figure S12M) and GATA3+ cells (Figure S12N), components of the type 2 immune response, were increased in both succinate-treated and AtohKO tissues (Schneider *et al*, 2018; Lei *et al*, 2018; Nadjsombati *et al*, 2018; Allen & Sutherland, 2014; Yamashita *et al*, 2004). These results demonstrate that commensal-derived succinate is capable of inducing tuft cell hyperplasia, even in the absence of helminth infection or eukaryotic colonization (Figure S12O).

As the microbiome was necessary for *in vivo* tuft cell hyperplasia, we used sequencing of the V4 region in the 16s rRNA gene to assess changes in microbiome distribution in the AtohKO model (Goodrich *et al*, 2014). DNA extraction and 16s sequencing were performed from the ileal luminal contents of co-housed wildtype and AtohKO littermates. Principal coordinate analysis of beta-diversity measures, including Bray-Curtis (Figures S13A, S13D), unweighted UniFrac (Figures S13B, S13E), and weighted UniFrac (Figures S13C, S13F) indices, which measure inter-sample relatedness (Goodrich *et al*, 2014), demonstrated that wildtype and AtohKO replicates clustered together based on biological phenotype rather than due to cage effects. Analysis of microbiome composition revealed a decrease in genus *Barnesiella* within the AtohKO replicates compared to the wildtype replicates, while the relative abundance of *Parasutterella* and *Bifidobacterium* was increased (Figures S13G-S13I). The latter has been associated with ameliorating symptoms in a variety of gastrointestinal pathologies, including colorectal cancer and IBD (O’Callaghan & van Sinderen, 2016). *Bifidobacterium infantis, Bifidobacterium breve,* and *Bifidobacterium pseudolongum* are components of the VSL3 probiotic, which has demonstrated the ability to induce remission in a subset of patients with active IBD (Bibiloni *et al*, 2005). We applied PiCRUST (Langille *et al*, 2013) to our 16s data and identified eight functional categories that were positively enriched in the AtohKO microbiome, including “Chlorocyclohexane and chlorobenzene degradation” and “Retinol metabolism” (Figure 4N) (Langille *et al*, 2013). Investigation of the Chlorocylohexane and chlorobenzene degradation category (ko00361) revealed that this pathway, simplified in the supplement, was associated with succinate production (Figure S13J). Further analysis revealed that *Bifidobacterium*, *Lactobacillus*, *Sutterellla*, *Acinetobacter*, and *Akkermansia* contributed to the positive enrichment of this pathway in the AtohKO microbiome (Figure 4O), however only *Bifidobacterium* was significantly increased compared to the wildtype small intestine (Figure 4O). Specifically, we observed that *Bifidobacterium pseudolongum*, a known producer of succinic acid (Van der Meulen *et al*, 2006), was increased six-fold in the AtohKO microbiome (Figure S13K). Our findings suggest that, in the absence of eukaryotic colonization, the commensal microbiome produces succinate and drives tuft cell hyperplasia, potentially via the type 2 immune response.

### Succinate treatment ameliorates inflammation in the TNF^ΔARE/+^ model

Metabolic and microbiome analysis of the AtohKO small intestine indicated that the TCA cycle intermediate succinate was linked to the *in vivo* tuft cell hyperplasia phenotype. Given the inverse correlation between tuft cell frequency and inflammation severity in the human and mouse ilea, we hypothesized that enhanced tuft cell specification could suppress inflammatory disease. Therefore, we therapeutically administered succinate in the drinking water of adult TNF^ΔARE/+^ mice, following disease onset, for a short-(<1 week) and long-term (1 month+) treatment period. While succinate-treated TNF^ΔARE/+^ animals failed to gain as much weight as untreated controls, this can likely be attributed to the palatability of succinate, as wildtype animals treated with succinate also had minimal weight gain (Figure S14A). However, succinate treatment in TNF^ΔARE/+^ animals markedly improved intestinal tissue organization compared to age-matched, untreated TNF^ΔARE/+^ controls, based on restored crypt-villus architecture and minimized villus distortion (Figure 5A). Histological examination demonstrated that succinate-treated animals had decreased mucosal destruction, as assessed by depth of inflammation and extent of tissue injury throughout the length of the ileum (Figure 5B-C). Additionally, we assessed immune cell subsets, including neutrophils and T-regulatory cells, that are increased in the TNF^ΔARE/+^ model (Kontoyiannis *et al*, 2002; Chang *et al*, 2012). MPO+ neutrophils (Figure 5F) and FOXP3+ T-regulatory cells (Figure 5G) were significantly reduced in succinate-treated animals, indicative of decreased infiltrative disease. In order to determine alterations in the type 2 immune response following succinate treatment, we examined the presence of GATA3-positive lymphocytes in untreated TNF^ΔARE/+^ animals (Yamashita *et al*, 2004). We observed high GATA3 expression within inflammatory infiltrates of the submucosal layer but closer examination revealed that the GATA3 signal did not co-localize with CD3e (Figure S14B i), indicating that they may be from non-lymphocyte-derived sources. Furthermore, GATA3 was also detected in the muscle layers (Figure S14B ii), possibly due to the increased presence of inflammation-activated stromal elements, such as intestinal myofibroblasts that are known to activate this transcription factor under inflammatory conditions (Roulis *et al*, 2011; Kimura *et al*, 2006). However, in the lamina propria of these animals, GATA3 signal was lower than in other regions and co-localization of CD3e and GATA3 remained sparse (Figure S14B), consistent with previous work in this model which delineated a role for Th1/Th17 immunity in driving inflammation (Kontoyiannis *et al*, 2002, 1999; Roulis *et al*, 2011). Short-term succinate treatment enhanced the presence of lamina propria CD3e+ GATA3+ cells in regions of resolved inflammation (Figure S14C). In long-term succinate treated animals, these cells were absent in the intervillus compartments, following resolution of inflammation (Figure S14D). Decreased GATA3+ stromal elements were observed in short-term treated animals, and these elements were mostly ablated following long-term treatment, restoring wildtype architecture (Figure S14D). This suggests that the Type 2 immune response, which is known to be activated following succinate treatment, accompanies the dampening of inflammation in the TNF^ΔARE/+^ model.

**Figure 5.**
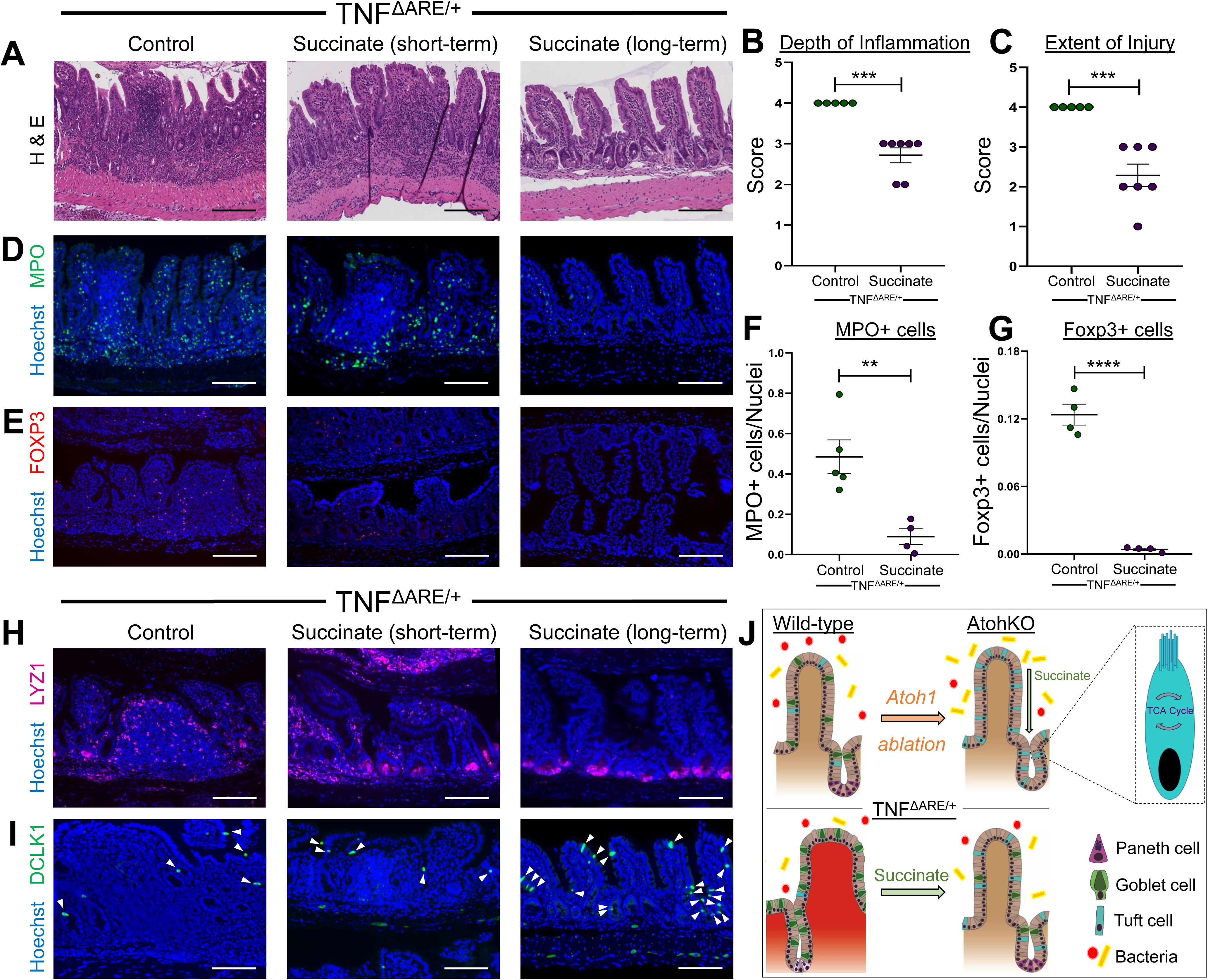
Therapeutic succinate treatment mitigates inflammation in the TNF^ΔARE/+^ model. **(A)** Histology from control and succinate-treated (120 mM) TNFΔARE/+ ileum. Scale bar = 100µm. **(B-C)** Pathological scoring of **(B)** depth of inflammation (0-4) and **(C)** tissue injury (0-4) in control and succinate-treated TNFΔARE/+ mice. Error bars represent SEM from n = 5 control and n = 7 succinate-treated TNFΔARE/+ mice. ***p-value < 0.001 by t-test. **(D-E)** Immunofluorescence imaging of **(D)** MPO (green) and **(E)** FOXP3 (red) in the ileum of untreated and succinate-treated TNFΔARE/+ ileum. Hoechst (blue) denotes nuclei, scale bar = 100 µm. **(F-G)** Quantification of **(F)** MPO+ and **(G)** FOXP3+ cells normalized to nuclei number. Error bars represent SEM from n = 4-5 untreated and n = 4 succinate-treated mice. **p-value < 0.01 and ****p – value < 0.0001. **(H-I)** Immunofluorescence imaging of (H) LYZ1 (magenta) and **(I)** DCLK1 (green) in the ileal epithelium of untreated and succinate-treated TNFΔARE/+ ileum. Scale bar = 100 µm. White arrows indicate DCLK1+ epithelial tuft cells and Hoechst (blue) denotes nuclei. **(J)** Summary diagram of findings.

Furthermore, we observed that markers of epithelial organization, including expression of epithelial LYZ1, increased in short-term treated animals and appeared to be completely restored with long-term treatment (Figure 5H). Strikingly, LYZ1 expressing cells were restricted to the base of the epithelial crypts and exhibited the typical Paneth cell morphology, while stromal LYZ1 expression was absent, resembling the wildtype ileal mucosa (Figure 5H). As hypothesized, DCLK1+ tuft cells were increased in long term-treated animals, consistent with succinate’s ability to drive tuft cell specification (Figure 5I). scRNA-seq analysis revealed that the canonical succinate receptor gene *Sucnr1* was highly enriched in wildtype ileal tuft cells compared to non-tuft cell populations (Figure S14E), further suggesting that the actions of succinate in suppressing inflammation were mediated through epithelial tuft cells rather than other cell types (Nadjsombati *et al*, 2018; Connors *et al*, 2018). Previous work demonstrated differences in tuft cell specification between the small and large intestine, with the majority of tuft cell studies focusing on the small intestine population (Herring *et al*, 2018; McKinley *et al*, 2017; Gerbe & Jay, 2016). scRNA-seq analysis of the colonic epithelium (Figures S14F-S14G) demonstrated that colonic tuft cells lack *Sucnr1* expression (Figure S14H). Consequently, neither DCLK1+ tuft cells (Figure S14I) nor eosinophils (Figure S14J) were increased in the colon following succinate treatment. Overall, these results implicate a role for enhanced tuft cell specification in suppressing ileal inflammation in the TNF^ΔARE/+^ model (Figure 5I).

## Discussion

Recent publications have provided significant insight into the sentinel role of tuft cells during acute helminth infection in the gastrointestinal tract (Gerbe & Jay, 2016; Gerbe *et al*, 2016). Chemosensation of helminth-derived succinate enables tuft cells to detect pathogen invasion, activate a type 2 immune response, and, subsequently, induce parasite expulsion (Nadjsombati *et al*, 2018; Lei *et al*, 2018). Tuft cell misspecification results in a dampened type 2 immune response and sustained worm burden, highlighting the necessity for this rare cell type in host response to helminth infection (Schneider *et al*, 2018). While endemic helminth infection is a global health concern, helminth therapy has also been proposed for the treatment of CD (Broadhurst *et al*, 2010; Helmby, 2015; Summers *et al*, 2005a). Prevailing thought postulates that the immune response mounted as a consequence of helminth infection may counteract the pro-inflammatory signaling driving CD (Summers *et al*, 2005a). This study presents the first evidence suggesting that small intestinal tuft cells possess the capacity to modulate intestinal inflammation. We observed decreased numbers of tuft cells, labeled with pEGFR and COX2, in ileal tissues acquired from CD patients. These results were confirmed in the TNF^ΔARE/+^ model of Crohn’s-like ileitis where frequency of DCLK1+ tuft cells was inversely correlated with inflammation severity. We hypothesized that tuft cell presence may act to suppress inflammation and, conversely, increasing tuft cell specification may potentially counteract pro-inflammatory signals in the inflamed intestine. To identify mechanisms driving increased specification of tuft cells, we developed a genetically-inducible model of tuft cell hyperplasia, the AtohKO model. We performed scRNA-seq in wildtype, TNF^ΔARE/+^, and AtohKO animals to examine gene expression changes in small intestinal tuft cells. Analysis of wildtype and TNF^ΔARE/+^ scRNA-seq datasets using the p-Creode algorithm confirmed a non-secretory origin for epithelial tuft cells, independent of the *Atoh1*-dependent secretory lineage (Gracz *et al*, 2018). In the AtohKO model, we applied both trend dynamic and population analysis to demonstrate that expression of TCA cycle genes was upregulated. This suggests that tuft cells in the AtohKO model are specified in a more metabolically favorable environment.

Given the chemosensory role associated with tuft cells, we asked whether these metabolic changes arose from non-cell autonomous mechanisms. Functional analysis of changes in the AtohKO microbiome implicated the production of the TCA cycle intermediate, succinate. Increased levels of *Bifidobacterium* and succinate were detected in the AtohKO intestine, suggesting a correlation to the tuft cell hyperplasia observed. Several strains of the genus *Bifidobacterium* are known succinic acid producers and have been linked to establishment of an adult microbiome during infant development as well as favorable outcomes in colon health (Van der Meulen *et al*, 2006; O’Callaghan & van Sinderen, 2016). Reduced levels of *Bifidobacterium* are associated with CD dysbiosis and numerous *Bifidobacterium* species are constituents of the VSL3 probiotic, indicating that the genus is necessary for maintenance of healthy intestinal function (Gerritsen *et al*, 2011; Bibiloni *et al*, 2005). As microbiome depletion in the AtohKO model suppressed tuft cell hyperplasia and decreased expression of TCA cycle genes, we hypothesize that the microbiome, based on the enrichment of succinic acid-producing species, may play a role in driving tuft cell specification in the AtohKO model. While parasite-derived succinate is known to signal to tuft cells in acute infection models, our findings demonstrate that commensal-derived succinate may also induce tuft cell hyperplasia.

In order to investigate the effect of increased tuft cell specification on inflammation, we therapeutically administered succinate to TNF^ΔARE/+^ animals following onset of disease. We observed significant histological improvements in succinate-treated TNF^ΔARE/+^ animals, including decreased villus blunting and tissue destruction compared to untreated controls. Immune cell infiltration, a hallmark of TNF^ΔARE/+^ ileitis, was significantly reduced based on the absence of MPO+ neutrophils and FOXP3+ T-regulatory cells. Finally, LYZ1+ Paneth cells were restored to the crypts of the ileum, comparable to wildtype controls, and less stromal LYZ1 expression was detected. Furthermore, DCLK1+ tuft cells were increased with succinate treatment in less inflamed TNF^ΔARE/+^ subjects, once more suggesting a role for this cell type in suppressing inflammation. Tuft cells sit at the crossroads of the epithelial, microbiome, and immune systems of the intestinal tract and, therefore, may have the potential to shift the balance between health and disease. Our findings demonstrate that enhanced tuft cell specification could be a viable therapeutic strategy for the treatment of inflammatory illnesses. Further studies exploring the mechanisms linking tuft cell specification to inflammatory signaling could lead to the development of clinically-viable strategies for the treatment of CD.

While numerous groups have demonstrated a role for succinate in activating type 2 immunity, it is unclear what mechanisms link succinate sensation with cytokine gene expression and immune cell activation. However, substantial evidence has demonstrated the ability of microbial-derived metabolites, to modulate gastrointestinal health, often by reprogramming of the host epigenome (Mathewson *et al*, 2016; Sharon *et al*, 2014; Cortese *et al*, 2016; Qin & Wade, 2018). Butyrate, a short-chain fatty acid (SCFA), produced via bacterial fermentation of dietary fermentation, promotes maturation of colonic T-regulatory cells (Smith *et al*, 2013; Arpaia *et al*, 2013) and is preferentially utilized as an energy source for epithelial colonocytes (Kaiko *et al*, 2016; Mathewson *et al*, 2016). A known histone deacetylase inhibitor, butyrate represses growth in proliferative progenitors and stem cells in the base of the colonic crypt (Kaiko *et al*, 2016; Kaiko & Stappenbeck, 2014). Folate, or vitamin B9, is essential for human health and is involved in DNA replication and nucleotide synthesis (Strozzi & Mogna, 2008; den Besten *et al*, 2013). While mammals cannot produce folate, certain *Bifidobacterium* and *Lactobacillus* species can generate this micronutrient from dietary sources (Strozzi & Mogna, 2008). Human subjects colonized with *Bifidobacterium* and *Lactobacillus* had increased folate levels as well as altered DNA methylation patterns in intestinal epithelial cells, suggesting that exposure to microbial signals can induce transcriptional changes in the host (den Besten *et al*, 2013; Cortese *et al*, 2016). Additionally, TCA cycle metabolites have been shown to induce epigenetic changes under favorable metabolic circumstances. For instance, the TCA cycle intermediate α-ketoglutarate is a substrate for a family of demethylase enzymes (Salminen *et al*, 2014), while Acetyl-CoA, in nutrient-rich conditions, induces cell growth by directly promoting histone acetylation (Mews *et al*, 2017; Cai *et al*, 2011). Therefore, it is possible that succinate acts in a similar manner to alter transcriptional patterns in the intestinal epithelium and induce downstream immunological changes.

## Methods

### Human Tissue

Formalin-fixed, paraffin-embedded blocks of ileum surgical resections were obtained from the Vanderbilt Cooperative Human Tissue Network Western Division, along with deidentified patient data and pathology reports. All studies were performed according to Vanderbilt University Institutional Review Board No. 182138. Pathological examination was utilized to classify samples as “normal” (n = 14) or “diseased” (n = 19). Samples from patients with Crohn’s disease were included only if inflammation was evident in the distal ileum. Tissue samples were prepared for immunofluorescence imaging as described below.

### Mouse Experiments

All animal protocols were approved by the Vanderbilt University Animal Care and Use Committee and in accordance with NIH guidelines. *Lrig1^CreERT2^* and *Atoh1^flox/flox^* strains, each in a C57BL/6 background, were purchased from Jackson Laboratory to generate *Lrig1^CreERT2/+^;Atoh1^fl/fl^* (AtohKO) animals. Cre recombinase activity was induced in two- or three-month-old *Lrig1^CreERT2/+^;Atoh1^fl/fl^* and *Lrig1^+/+^;Atoh1^fl/fl^* males via intraperitoneal administration of 2mg of tamoxifen (Sigma-Aldrich) for four consecutive days. Mice were sacrificed and tissues were harvested twenty-one days after the first injection. TNF^ΔARE/+^ and wildtype littermates were sacrificed at four months of age after disease onset, followed by tissue collection. Animal weights were recorded at regular intervals.

For microbiome depletion experiments, *Lrig1^CreERT2/+^;Atoh1^fl/fl^* animals were pre-treated with a broad-spectrum antibiotic cocktail containing kanamycin (4.0 mg/ml), metronidazole (2.15 mg/ml), gentamicin (0.35 mg/ml), colistin sulfate (8500 U/ml), and vancomycin (0.45 mg/ml) or ampicillin (1 mg/ml) in their drinking water for seven days prior to tamoxifen treatment. Mid-dose antibiotics and low-dose antibiotics were 0.75x and 0.25x of the original 1x concentration, respectively.

Following tamoxifen administration, *Lrig1^CreERT2/+^;Atoh1^fl/fl^* received either standard or antibiotic-supplemented drinking water for an additional fourteen days. For microbiome experiments, co-housed male *Lrig1^CreERT2/+^;Atoh1^fl/fl^* littermates received either vehicle (corn oil) or tamoxifen for 1 month. For succinate treatment, TNF^ΔARE/+^ received either sodium succinate hexahydrate (120 mM; Alfa Aesar) or standard drinking water following disease onset (three-to four-months-old) for five days or one month (Lei *et al*, 2018; Schneider *et al*, 2018).

### Immunofluorescence staining and imaging

Paraffin-embedded ileal tissues were section (5 µm) prior to deparaffinization, rehydration, and antigen retrieval using a citrate buffer (pH 6.0) for 20 minutes in a pressure cooker at 105°C, followed by a 20-minute cool down at room temperature (RT). Endogenous background signal was quenched by incubating tissue slides in 3% hydrogen peroxide for 10 minutes at RT. Tissue sections were blocked in staining buffer (3% bovine serum albumin/ 10% normal donkey serum) for 1 hour at RT prior to incubation with primary antibody overnight at RT. Antibodies used for immunofluorescence included those against LYZ1 (DAKO, 1:100, rabbit), MUC2 (Santa Cruz, 1:100, rabbit), MPO (DAKO, 1:100, rabbit), DCLK1 (Santa Cruz, 1:100, goat), FOXP3 (eBioscience, 1:50, rat), and GATA3 (Abcam, 1:50, rabbit). Sections were then incubated with AlexaFluor (AF)-555- or AF-647-conjugated secondary antibodies (Life Technologies, 1:500) for 1 hour at RT and Hoechst (1:10,000, Life Technologies) for 10 min at RT. For human tuft cell labeling, AF-488-conjugated pEGFR (Abcam, 1:100) and unconjugated COX2 (CST, 1:100, rabbit) were used. Slides were incubated with anti-rabbit AF-647-conjugated secondary antibody and stained with Hoechst. Imaging was performed using a Zeiss Axio Imager M2 microscope with Axiovision digital imaging system (Zeiss, Jena GmBH, Germany).

### Image quantification

To quantify human tuft cells, the number of crypt and villus structures were counted per field of view (FOV) for each subject (approximately 15-20 FOVs per sample). Tuft cell number, as identified by pEGFR and COX co-labeling, was manually counted per FOV in a blinded fashion. Quantification of tuft cell number per epithelial structure (crypt and villus) was generated for each subject and then stratified by disease state (“normal” or “Crohn’s disease”) based on the pathology report. Results were analyzed by t-test using Prism GraphPad. For MUC2 quantification, manual demarcation of the epithelial villi was performed in each FOV and a nuclear mask was generated based on Hoechst staining. In the same region, a MUC2 mask was generated using immunofluorescence staining of MUC2. Total area of both masks was calculated to generate a normalized ratio of MUC2 intensity to nuclear staining. This process was repeated for LYZ1 quantification, except that only crypts were manually demarcated in the FOVs. For MPO and DCLK1 quantification, each villus was considered a separate unit and number of MPO+ neutrophils and DCLK1+ tuft cells were counted in a blinded fashion. Tuft cell number per villi was stratified based on MPO staining as either “low inflammation” (<35 neutrophils per villi) or “high inflammation” (≥ 35 neutrophils per villi). Total number of villi counted, and total number used for significance testing by t-test can be found in Figure S2. Finally, for FOXP3 and MPO quantification, total area of each mask was used to calculate a normalized ratio of signal to nuclear staining.

### Immunohistochemistry and histological scoring

Paraffin-embedded ileal tissue from wildtype and succinate-treated slides were retrieved as described above and standard H&E staining was performed for histology. To identify eosinophils, tissues were incubated with anti-MBP (Mayo Clinic Arizona) followed by anti-rat HRP and counterstained with hematoxylin. Immunohistochemistry was performed by the Vanderbilt Translational Pathology Shared Resource and 20x brightfield scanning of immunohistochemistry slides was performed by a Leica SCN400 Slide Scanner in the Vanderbilt Digital Histology Shared Resource. Degree of inflammation in the distal ileum was assessed in a blinded fashion by a trained pathologist. Damage to the epithelium was assessed using depth of inflammation on a five-point scale (0 = no damage, 1 = mucosal damage only, 2 = submucosal infiltration, 3 = infiltration into muscularis propria, and 4 = infiltration into the periintestinal fat), while extent of inflammatory injury in the tissue swiss roll was also scored on a five-point scale (0 = <10%, 1 = 10-25%, 2 = 25-50%, 3 = 50-75%, and 4 = >75%).

### Enteroid Experiment

Ileal tissue was dissected and incubated in chelation buffer (3mM EDTA/EGTA, 0.5mM DTT, 1% P/S) at 4°C for 45 minutes. The tissue was shaken in PBS and filtered through a 100µm filter to isolate individual ileal crypts. The crypt suspension was centrifuged at 2.8 x 1000 RPM for 1 ½ minutes at 4°C following which 10µl of crypt pellet was resuspended in 300µl of reduced growth factor Matrigel and embedded in a 24-well dish. Enteroids were cultured initially in IntestiCult Organoid Growth Medium (StemCell Technologies) supplemented with Primocin antimicrobial reagent (InvivoGen, 1:1000) for 4 days before being changed to Primocin-supplemented differentiation media, as previously described (Sato *et al*, 2009). For *ex vivo* Cre-activation, *Lrig1^CreERT2/+^;Atoh1^fl/fl^* enteroids were treated overnight at 37°C with either 1 µM 4-hydroxytamoxifen (Sigma-Aldrich) or vehicle (ethanol) in differentiation media. Enteroids were passaged the following day into 8-well chamber slides and, after 5 days, were fixed on ice using 4% PFA for immunofluorescence staining.

### Enteroid Immunofluorescence Staining

Fixed enteroids were permeabilized with Triton X-100 for 30min and blocked with 1% normal donkey serum (PBS) for 30min at RT. Enteroids were stained with primary antibodies against LYZ1 (DAKO, 1:100, rabbit), MUC2 (Santa Cruz, 1:100, rabbit), and DCLK1 (Santa Cruz, 1:100, goat). Enteroids were then incubated with AF-555- or AF-647-conjugated secondary antibodies (Life Technologies, 1:500) for and Hoechst (1:10,000, Life Technologies) 1 hour at RT. Vectashield Antifade Mounting Medium (Vectorlabs) was applied to enteroids before imaging with a Nikon Spinning Disk Confocal microscope.

### Droplet-based single-cell RNA-sequencing

Ileal crypts from human and mouse tissue were isolated as described above. Crypts were dissociated into single cells using a cold-activated protease (1 mg/ml)/ DNAseI (2.5 mg/ml) enzymatic cocktail in a modified protocol that maintains high cell viability (Adam *et al*, 2017). Dissociation was performed at 4°C for 15 minutes followed by trituration to mechanically disaggregate cell clusters. Cell viability was assessed by counting Trypan Blue positive cells. The cell suspension was enriched for live cells with a MACS dead cell removal kit (Miltenyi) prior to encapsulation. Single cells were encapsulated and barcoded using the inDrop platform (1CellBio) with an *in vitro* transcription library preparation protocol (Klein *et al*, 2015). Briefly, the CEL-Seq work flow entailed (1) reverse transcription (RT), (2) ExoI digestion, (3) SPRI purification (SPRIP), (4) Second strand synthesis, (5) T7 *in vitro* transcription linear Amplification, (7) SPRIP, (8) RNA Fragmentation, (9) SPRIP, (10) primer ligation, (11) RT, and (12) library enrichment PCR (Herring *et al*, 2018). Each sample was estimated to contain approximately 2,500 encapsulated cells.

Following library preparation, the samples were sequenced using Nextseq 500 (Illumina) using a 150bp paired-end sequencing kit in a customized sequencing run (Herring *et al*, 2018). After sequencing, reads were filtered, sorted by their barcode of origin, and aligned to the reference transcriptome using the inDrop pipeline. Mapped reads were quantified into UMI-filtered counts per gene, and barcodes that corresponded to cells were retrieved based on previously established methods (Klein *et al*, 2015). Overall, from approximately 2,500 encapsulated cells, approximately 1,800-2,000 cells were retrieved per sample.

### Pre-processing and batch correction of scRNA-seq data

Datasets were filtered for cells with low library size or high mitochondrial gene expression (Klein *et al*, 2015). Filtered datasets for each replicate were analyzed using the Seurat pipeline (Stuart *et al*, 2018). Briefly, count matrices were log scale normalized followed by feature selection of highly variable genes. Canonical correlation analysis (CCA) was used to align replicates based on biological condition using dynamic time warping. Following subspace alignment, modularity optimization (0.8 resolution) was used to identify cell clusters. The ComBat algorithm (Chen *et al*, 2011) was then used to batch correct each gene on a per cluster basis. Visual assessment of alignment between replicates was performed using t-SNE analysis (Van Der Maaten & Hinton, 2008).

### p-Creode mapping and trajectory analysis

The wildtype dataset, as well as the GSE92332 ileum dataset, were feature selected using the binned variance method and these features were used for all other conditions (Butler *et al*, 2018). Feature selected datasets were analyzed using the p-Creode algorithm (https://github.com/KenLauLab/pCreode) (Herring *et al* 2018). For graph scoring, 100 independent runs were generated from each combined dataset. Overlay of ArcSinh normalized expression data was used to identify cell lineages and quantify tuft cell placement as either “secretory” or “non-secretory.”

### Trend analysis overview

Trend analysis was performed to identify gene expression changes in the AtohKO tuft cell lineage compared to the wildtype tuft cell lineage. 10 p-Creode resampled runs were used from the wildtype and AtohKO dataset. The top 2,500 genes (ranked by variance) over the tuft cell trajectory were selected from each of the wildtype and AtohKO datasets. The union of these gene sets (3,420 genes) was used for downstream analysis. The dynamic trend of gene expression for each gene over the tuft cell trajectory was obtained by fitting a linear Generalized Additive Model (GAM) with a normal distribution and an identity link function using 10 splines (Servén & Brummitt, 2018). The fitted curves were then normalized between 0 and 1 for comparison between datasets. For each gene trend, classification of its dynamics was performed by calculating its dynamic time warping distances to 12 reference trends. These categories were then broadly combined into 5 classes: (1) upward, (2) upward transitory, (3) downward transitory, (4) downward, and (5) flat. For the coarse grain analysis, 3 trend classes were formed by combining (1) group 1 and 2 genes into “upward,” (2) group 3 and 4 genes into “downward,” and (3) flat. Trend classification was scored by consensus over 10 resampled runs (Herring *et al*, 2018) for both the (A) 5-trend and (B) 3-trend analysis. Genes with high consensus are those with a cumulative sum of 16 between the two classifications (for instance, a gene being grouped in the same trend in 8 out of 10 p-Creode replicates in the 5-trend analysis and in 8 out of 10 p-Creode replicates in the 3-trend analysis). This resulted in 2,004 high-consensus genes being used for downstream over-representation analysis. From the list of high consensus genes, we identified genes which switched from (A) wildtype group 4 to AtohKO group 1, 2, or 3, (B) wildtype group 3 to AtohKO group 1 or 2, or (C) wildtype group 2 to AtohKO group 1. Over-representation analysis, based on KEGG, Reactome pathway, and Wiki pathway datasets, of upregulated genes was performed in WebGestalt (http://www.webgestalt.org/) (Wang *et al*, 2017). Using 3-trend analysis, we identified genes that switched from wildtype group 2 (downward) to AtohKO group 1 (upward) and performed over-representation analysis.

### Visualization and significance testing of trend analysis

Visualization of trend dynamics was performed using Matlab software for enriched genes from the Reactome Pathway “Citric acid cycle (TCA cycle)” (https://reactome.org/content/detail/R-HSA-71403) gene list. Each gene plot includes raw expression data from each wildtype or AtohKO p-Creode replicate and the trend line as an average of raw expression data aligned by dynamic time warping across all ten resampled runs for each respective condition. For significance testing between wildtype and AtohKO trends, randomized classifications were generated for each gene in the wildtype and AtohKO condition by resampling from the same distribution for each condition. The null hypothesis stated that there was no consensus across the wildtype classifications or that there was no upward class switching from wildtype to AtohKO trajectories. Simulations comparing the randomized and observed classifications were performed 10,000 times to obtain a p-value.

### Gene set enrichment analysis (GSEA) of differential expression

Median difference in gene expression was calculated between wildtype and AtohKO tuft cells. GSEA (http://software.broadinstitute.org/gsea/index.jsp) of differential gene expression was performed to identify positively enriched pathways. Top twenty gene sets with the highest NES and most significant p-value were used for the further analysis. Significance testing by t-test was used to compare relative expression of highly enriched genes between wildtype and AtohKO tuft cells using Prism Graphpad. GSEA and over-representation analysis for specific gene sets (KEGG, Reactome pathway, and Wiki pathways) was performed using WebGestalt (http://www.webgestalt.org/) (Wang *et al*, 2017).

### DNA extraction and 16s rRNA sequencing

Ileal luminal contents were collected fresh from tamoxifen- or vehicle-treated *Lrig1^CreERT2/+^;Atoh1^fl/fl^* animals, as described above. All samples were collected on the same day and frozen in 2ml Eppendorf tubes (DNAse and RNAse free) at −80°C. Microbial genomic DNA extraction was performed using the PowerSoil DNA Isolation Kit (MO BIO Laboratories) (Zackular *et al*, 2016). Briefly, luminal contents were added to PowerBead Tubes and homogenized twice in a Bead Beater machine for 3 minutes with a 2-minute cool down in between homogenization. Samples were then processed as per the kit instructions and DNA was eluted into a sterile buffer. The V4 region of the 16s rRNA gene from each sample was amplified and sequenced by Georgia Genomics and Bioinformatics Core (http://dna.uga.edu) using the Illumina MiSeq Personal Sequencing platform (Zackular *et al*, 2014).

Raw 16S rRNA sequences were filtered for quality (target error rate < 0.5%) and length (minimum final length 225bp) using Trimmomatic (Bolger *et al*, 2014) and QIIME (Caporaso *et al*, 2010). Spurious hits to the PhiX control genome were identified using BLASTN and removed. Passing sequences were trimmed of forward and reverse primers, evaluated for chimeras with UCLUST (Edgar & Bateman, 2010) (de novo mode), and screened for mouse-associated contaminant using Bowtie2 followed by a more sensitive BLASTN search against the GreenGenes 16S rRNA database. Chloroplast and mitochondrial contaminants were detected and filtered using the RDP classifier with a confidence threshold of 80%. High-quality 16S rRNA sequences were assigned to a high-resolution taxonomic lineage using Resphera Insight (Drewes *et al*, 2017). Functional gene content was inferred using PICRUSt (Langille *et al*, 2013). Statistical analyses between groups was performed using R (v3.5; http://cran.r-project.org/). Heatmap clustering visualizations were generated using Microsoft Excel, selecting microbiome features with FDR adjusted p-values < 0.10, and log-transforming values for color scaling.

### Pathogen testing

Fresh fecal samples from non-co-housed animals (n = 5) and frozen at −20°C. Samples were shipped to IDEXX BioAnalytics (Columbia, Missouri) for pathogen PCR panel and helminth float testing (Figure S12O).

### O-benzylhydroxylamine derivatization of cecal contents and tissue

O-benzylhydroxylamine (O-BHA) derivatization of common tricarboxylic acid intermediates was performed from both cecal luminal contents and tissues (Tan *et al*, 2014) by the Vanderbilt Mass Spectrometry Service Laboratory. Briefly, luminal filtrate was mixed in MeOH/H_2_O with 0.1% Formic acid and added to Pyr ^13^C_3_(327 µg/ml), EDC, and O-BHA as described in the Sherwood protocol (Fensterheim *et al*, 2018). Tissue homogenates were processed similarly, and both were incubated at room temperature for 1 hour prior to extraction with ethyl acetate. 100 µl of luminal content sample (200 mg/ml) or tissue sample (250 mg/ml) was analyzed for specified metabolites and analyte response ratios were calculated using validated standards (Zhu *et al*, 2014). An aliquot of PBS was processed and reconstituted as a negative control.

## Acknowledgements

K.S.L., A.B. and H.Y.K. are funded by R01DK103831. K.S.L., A.J.S., A.N.S., M.A.R.S., Q.L. are funded by U2CCA233291. K.S.L. and Q.L. are funded by P50CA095103. C.A.H. is funded by a training grant from T32HD007502 and a pre-doctoral F31GM120940. B.C. is funded by a training grant from T32LM012412. E.A.S is funded by KL2TR002245. K.S.L., E.A.S., and M.K.W. are funded by P30DK058404. M.K.W. is funded by UM1CA183727. J.R.W. is the founder of Resphera Biosciences. The authors declare no competing financial interests. Microbiome sequencing was funded by the Vanderbilt Microbiome Initiative. The authors would like to thank Cherie’ Scurrah and Paige Vega in the Lau lab for support in mouse experiments, Seth Bordenstein, Andrew Brooks, Nicholas Markham, Oliver McDonald, Eliot McKinley, Robert Coffey, Izumi Kaji, James Goldenring, Eunyoung Choi, Joseph Roland, Damian Maseda, the Vanderbilt Digestive Disease Research Center, and the Vanderbilt Epithelial Biology Center for helpful discussions centered around the microbiome, tuft cells, and metabolism. The authors would also like to thank various cores and centers at Vanderbilt for making this work possible, including the Vanderbilt Mass Spectrometry Research Center, Cooperative Human Tissue Network, VANTAGE Genomics Core, the Translational Pathology Shared Resource, and the Digital Histology Shared Resource.

## Author Contributions

A.B. conceived the study, performed the experiments, analyzed the data, compiled the figures, and wrote the manuscript. H.Y.K. performed image quantification analysis. A.J.S., A.N.S. performed single-cell RNA-seq experiments. C.A.H., B.C., M.A.R.S., Q.L. performed single-cell data analysis. J.R.W. performed microbiome data analysis. E.A.S. acquired human patient tissues. M.K.W. acquired human patient tissues and assisted in pathologic analysis. K.S.L. conceived the study, performed data analysis, participated in manuscript writing, and supervised the research.

---

**Figure S1.**
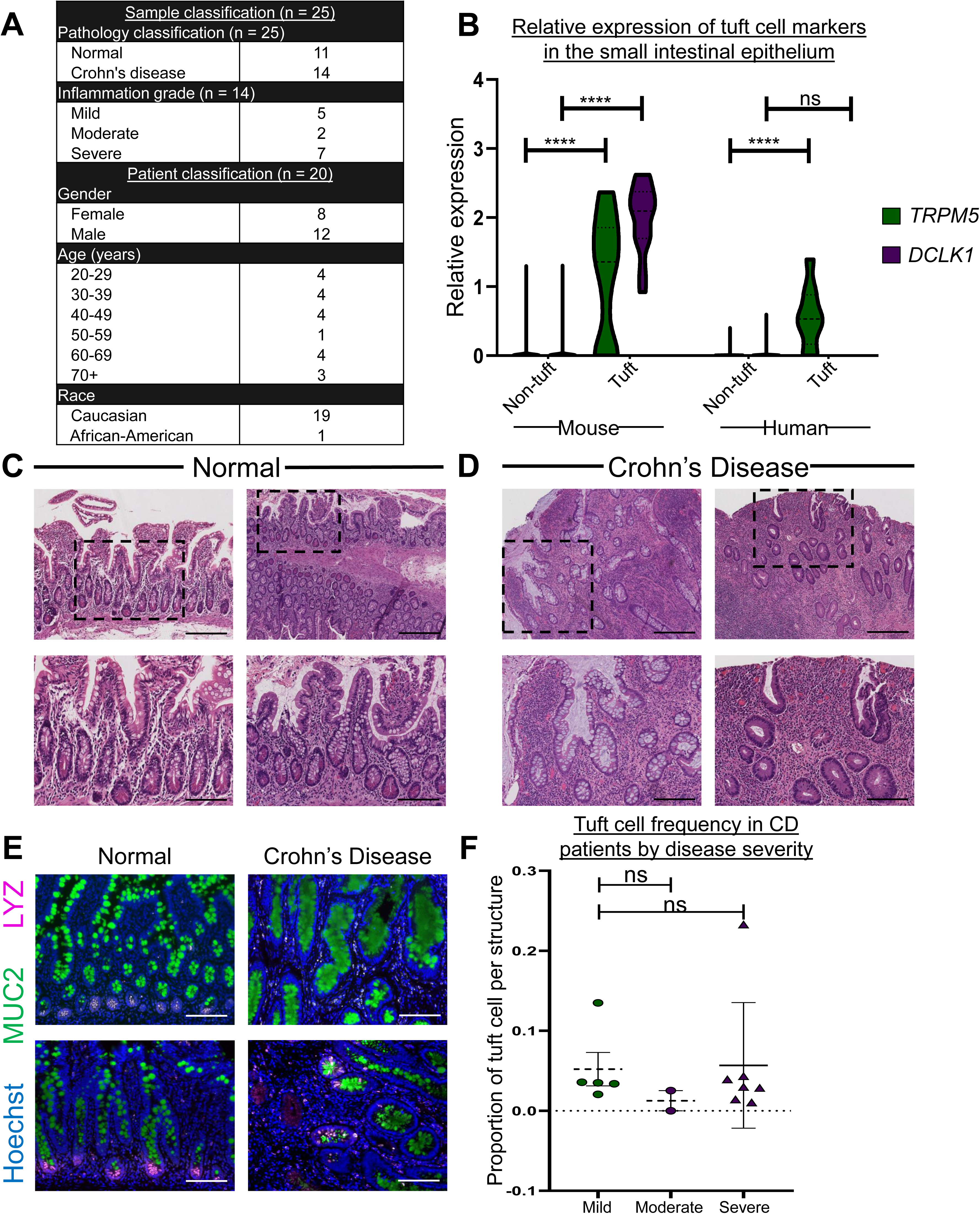
Human tuft cells are decreased in patients with ileal Crohn’s disease, Related to Figure 1. **(A)** Sample and patient summary statistics from ileal surgical resections. **(B)** Quantification of relative gene expression of *Dclk1* (green) and *Trpm5* (magenta) in mouse and human non-tuft (n = 1166 and 1942 cells, respectively) and tuft cell (n = 13 and 9 cells, respectively) populations from single-cell RNA sequencing. ****p-value < 0.0001 by t-test. N = 1 replicate per mouse and human**. (C-D)** Histology of the distal small intestine from **(C)** normal and **(D)** Crohn’s disease patients. Dotted square shows magnified inset. Representative images are shown from two separate normal or Crohn’s disease patients, respectively. Scale bar = 100 µm. **(E)** Immunofluorescence staining of MUC2 (green) and LYZ (magenta) in normal and Crohn’s disease ileal biopsies. Representative images are shown from two separate normal or Crohn’s disease patients, respectively. Hoechst (blue) denotes nuclei, scale bar = 100 µm. **(F)** Quantification of pEGFR and COX2 double-positive tuft cells in CD patients stratified by disease severity – mild (n = 5), moderate (n = 2), and severe (n = 7). Error bars present SEM and each dot represents a separate sample. Not significant (ns) by t-test.

**Figure S2.**
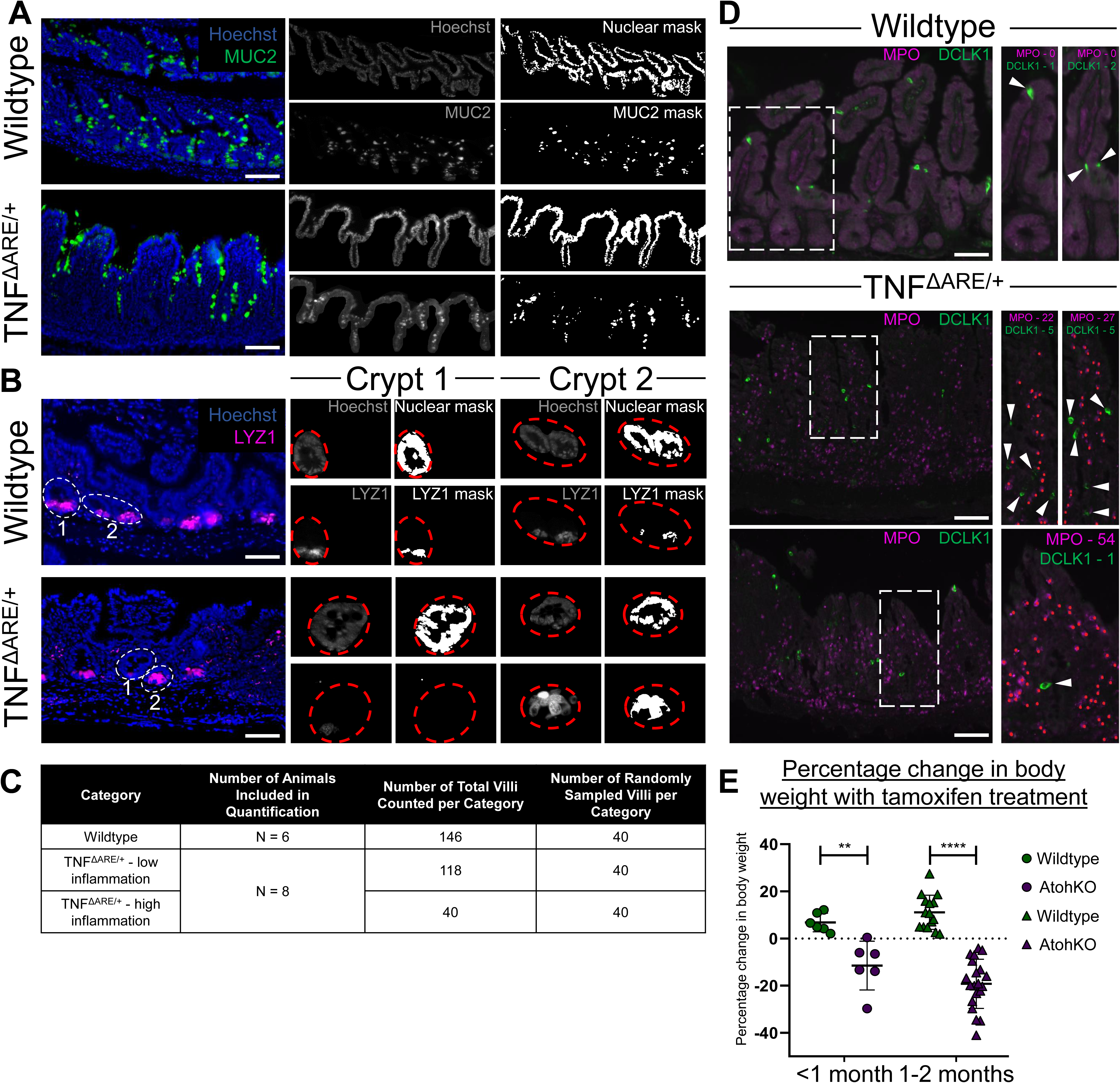
Image processing of immunofluorescence staining was used to quantify DCLK1+ tuft cell number versus severity of inflammation, Related to Figure 1. **(A)** Image analysis of Hoechst (blue) and MUC2 (green) staining were used to manually demarcate the epithelial monolayer and generate a nuclear and MUC2 mask, respectively, from wildtype, TNF^ΔARE/+^, and AtohKO (not pictured) images. MUC2 levels were normalized to total Hoechst area to quantify staining intensity between conditions. Scale bar = 100 µm. **(B)** Image analysis of Hoechst (blue) and LYZ1 (magenta) staining were used to manually demarcate individual crypts (white dashed line) and generate a nuclear and LYZ1 mask (red dashed line), respectively, from wildtype, TNF^ΔARE/+^, and AtohKO (not pictured) images. LYZ1 levels were normalized to total Hoechst area to quantity staining intensity. Scale bar = 100 µm. **(C)** Immunofluorescence imaging of MPO (magenta) and DCLK1 (green) was used to manually quantify severity of inflammation and tuft cell number, respectively. Within the magnified inset, white arrows indicate DCLK1+ tuft cells and red dots indicate MPO+ neutrophils. Scale bar = 100 µm. **(D)** Table 4 shows the total number of villi counted for DCLK1+ quantification. 40 villi were randomly selected for final quantification. **(E)** Percentage change in starting body weight following tamoxifen treatment in wildtype (green) and AtohKO (magenta) animals stratified by short-term (<1 month, circles) and long-term (1-2 months, triangles) treatment. Each data point represents a single animal and error bars represent SEM calculated from n = 6 <1 month wildtype and AtohKO each, n = 15 1-2 month wildtype, and n = 20 AtohKO animals. ****p-value < 0.0001, **p-value < 0.01 by t-test.

**Figure S3.**
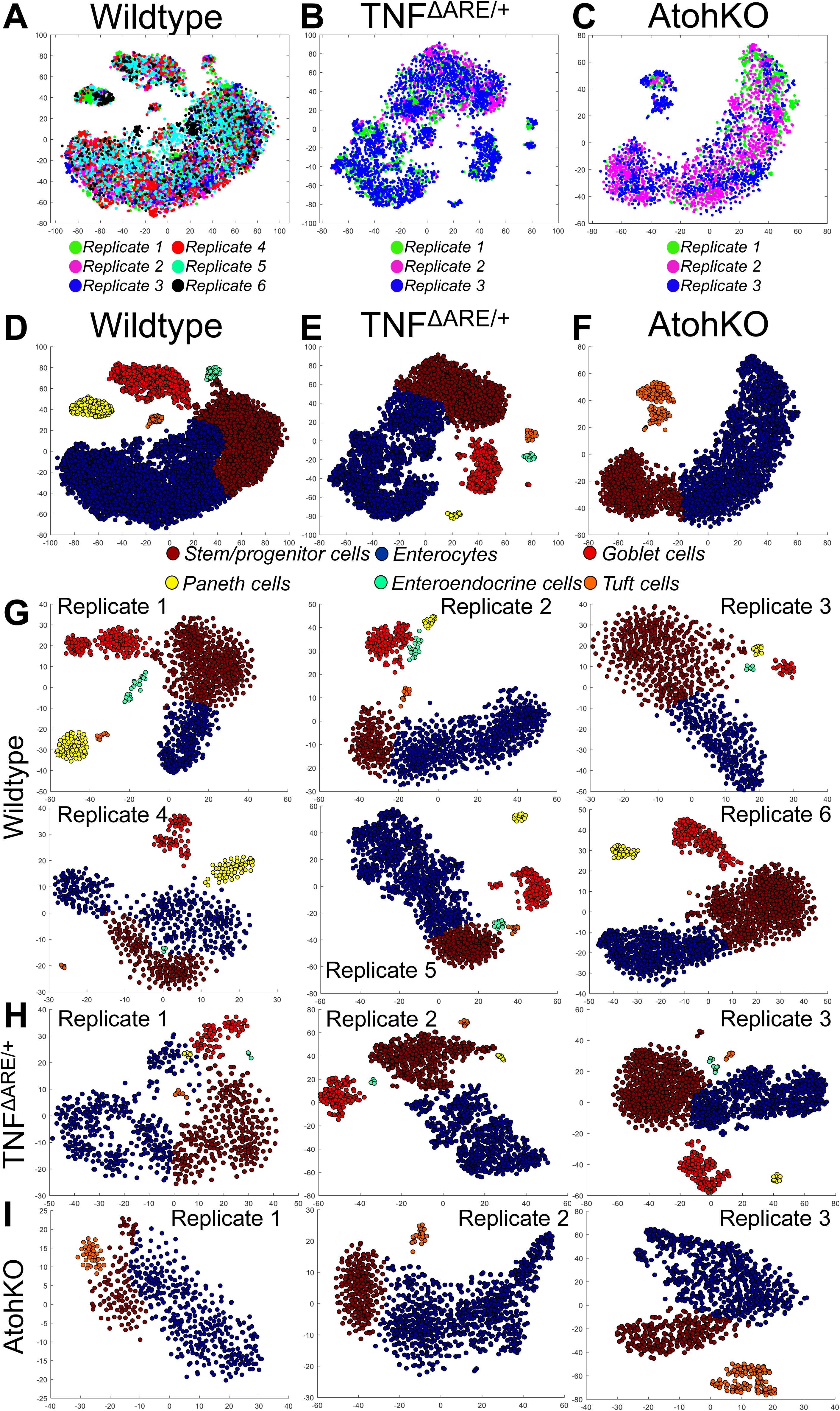
Batch alignment of scRNA-seq biological replicates, Related to Figure 2. **(A-C)** t-SNE analysis of the **(A)** wildtype (n = 6,932 cells). Manual annotation demarcates datapoints from individual replicates in the wildtype (n = 6 replicates) dataset (Rep. 1 – green, Rep. 2 – magenta, Rep. 3 – blue, Rep. 4 – red, Rep. 5 – teal, and Rep. 6 – black). Each datapoint represents a single cell. **(B-C)** TNF^ΔARE/+^ (n = 3,401 cells) and **(C)** AtohKO (n = 2,456 cells) scRNA-seq datasets. Manual annotation demarcates datapoints from individual replicates in the TNF^ΔARE/+^ (n = 3) and AtohKO (n = 3) datasets. Rep. 1 – green, Rep. 2 – magenta, and Rep. 3 – blue. Each datapoint represents a single cell. **(G-I)** Clustering analysis of replicates from the **(G)** wildtype, **(H)** TNF^ΔARE/+^, and **(I)** AtohKO scRNA-seq datasets to show stem/progenitor cells (brown), enterocytes (blue), goblet cells (red), Paneth cells (yellow), enteroendocrine cells (green), and tuft cells (orange).

**Figure S4.**
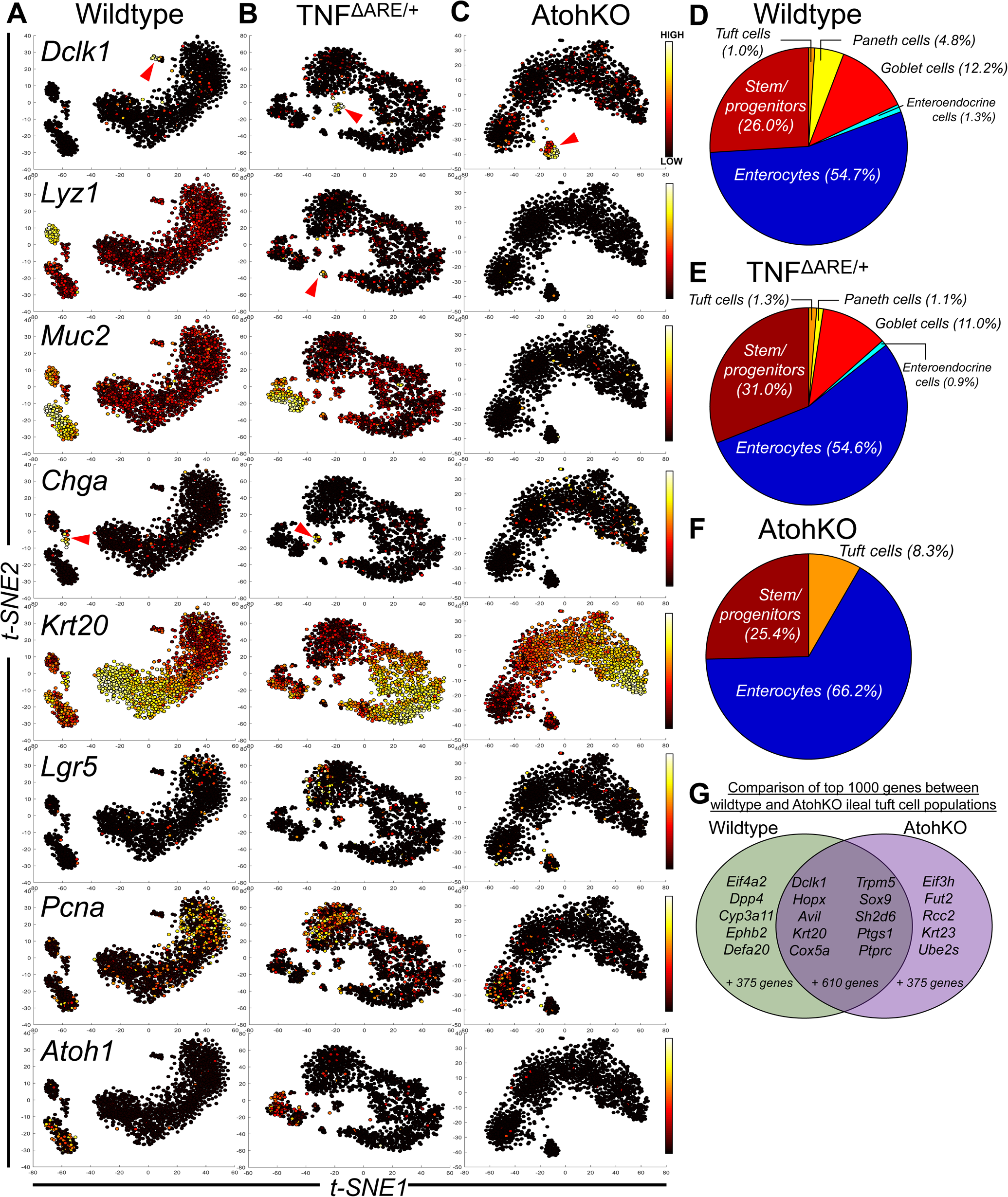
Distribution of epithelial cell populations in scRNA-seq datasets, Related to Figure 2. **(A-C)** t-SNE plots of scRNA-seq data generated from **(A)** wildtype, **(B)** TNF^ΔARE/+^, and **(C)** AtohKO ileal epithelium. Selected cell identity genes, including *Dclk1*, *Lyz1*, *Muc2*, *Chga*, *Krt20*, *Lgr5*, *Pcna*, and *Atoh1*, are overlaid on the t-SNE plots. Red arrows indicate smaller cell clusters, such as tuft and enteroendocrine populations. Each t-SNE plot depicts 1,450 randomly selected datapoints from their corresponding complete dataset and each datapoint represents a single cell. Overlay represents ArcSinh-scaled expression data. **(D-F)** Quantification of cell populations within the **(D)** wildtype, **(E)** TNF^ΔARE/+^, and **(F)** AtohKO scRNA-seq data. **(G)** Analysis of the 1,000 highest expressed genes in the wildtype (green) and AtohKO (magenta) tuft cell population. 620 genes are expressed in the union of both datasets while 380 genes are expressed in one but not the other dataset.

**Figure S5.**
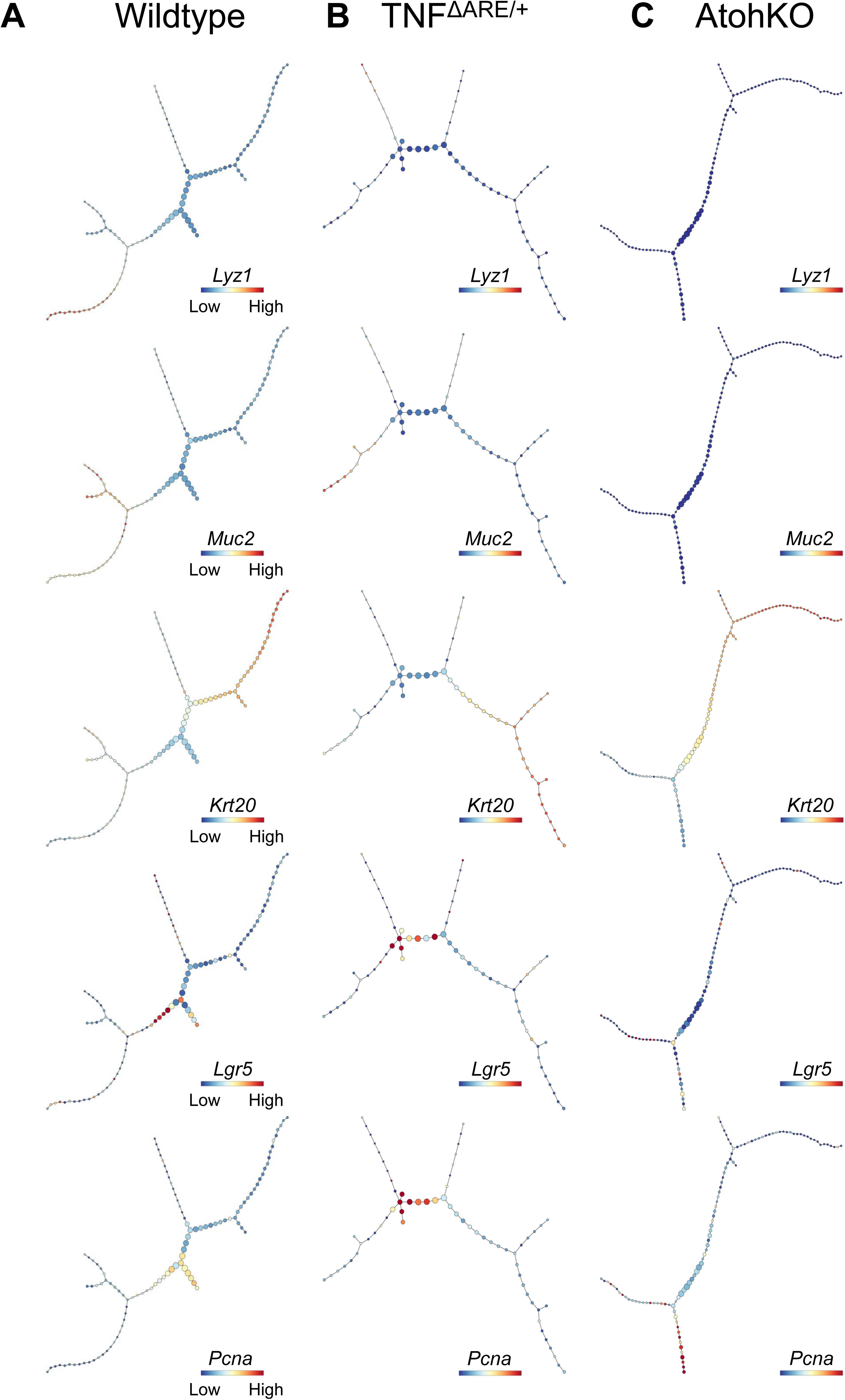
Gene expression overlay of epithelial cell identity genes on p-Creode graphs, Related to Figure 2. **(A-C)** Overlay of selected tuft cell-specific genes, including *Lyz1*, *Muc2*, *Krt20*, *Lgr5*, and *Pcna* on **(A)** wildtype, **(B)** TNF^ΔARE/+^, and **(C)** AtohKO p-Creode maps. Overlays represent ArcSinh-scaled gene expression data.

**Figure S6.**
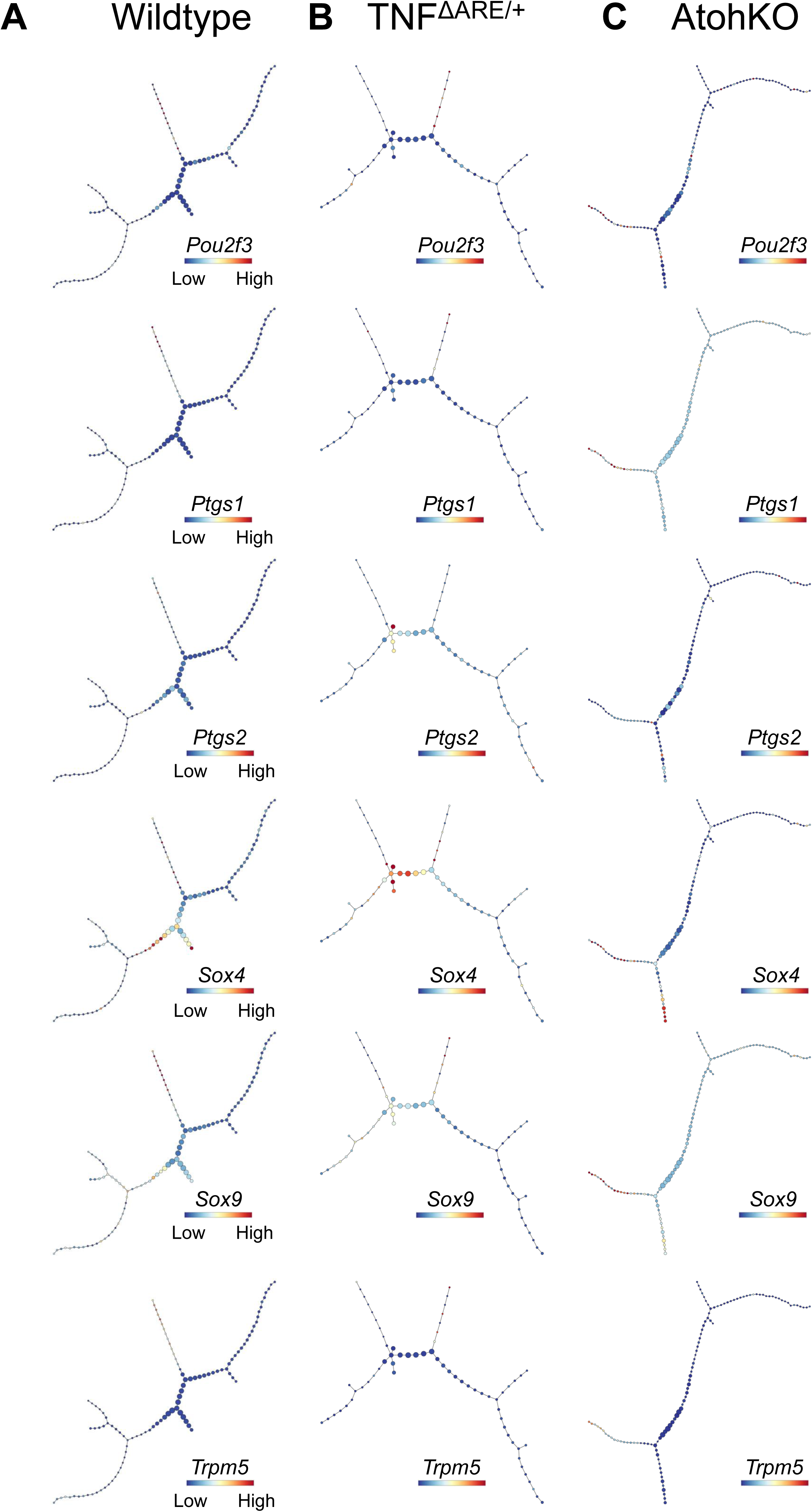
Gene expression overlay of tuft cell-specific genes on p-Creode graphs, Related to Figure 2. **(A-C)** Overlay of selected tuft cell-specific genes, including *Pou2f3*, *Ptgs1*, *Ptgs2*, *Sox4*, *Sox9*, and *Trpm5*, on **(A)** wildtype, **(B)** TNF^ΔARE/+^, and **(C)** AtohKO p-Creode maps. Overlays represent ArcSinh-scaled gene expression data.

**Figure S7.**
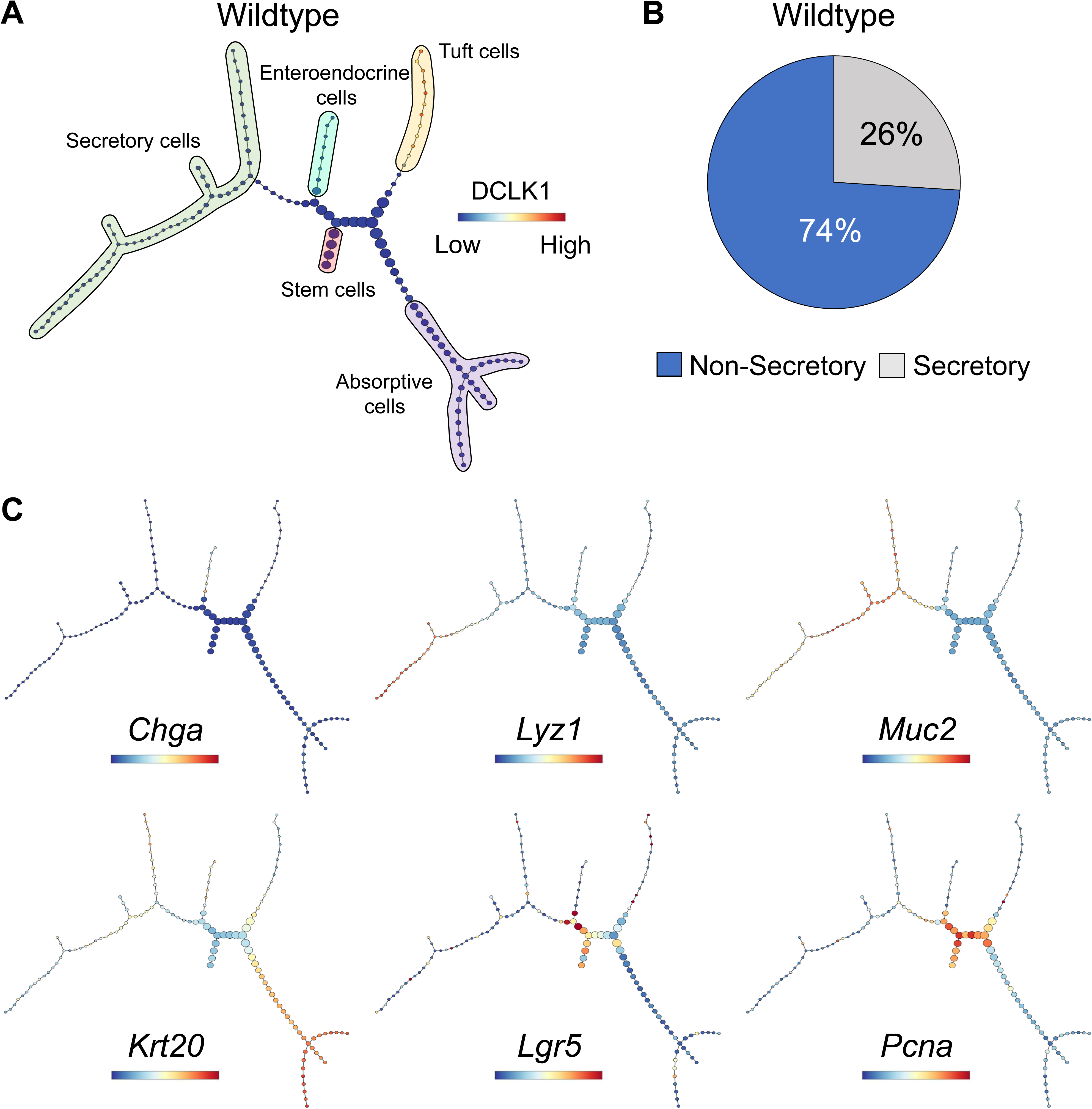
p-Creode analysis of small intestinal epithelium with enteroendocrine cells, Related to Figure 2. (A) Representative p-Creode map of wildtype scRNA-seq dataset, including enteroendocrine cells. Graph overlay depicts *Dclk1* levels. Cell lineages, including enteroendocrine cells (teal), secretory cells (green), absorptive cells (purple), tuft cells (orange), and stem cells (red), were manually labelled. Node size represents cell state density and each edge represents cell state transitions. Overlays represent ArcSinh-scaled gene expression data. (B) Quantification of wildtype maps (n = 100) indicating 74% of graphs were classified as non-secretory and 26% as secretory. (C) Overlay of selected cell identity genes, including *Chga*, *Lyz1*, *Muc2*, *Krt20*, *Lgr5*, and *Pcna*, depicting epithelial cell differentiation in the wildtype topology. Overlays represent ArcSinh-scaled gene expression data.

**Figure S8.**
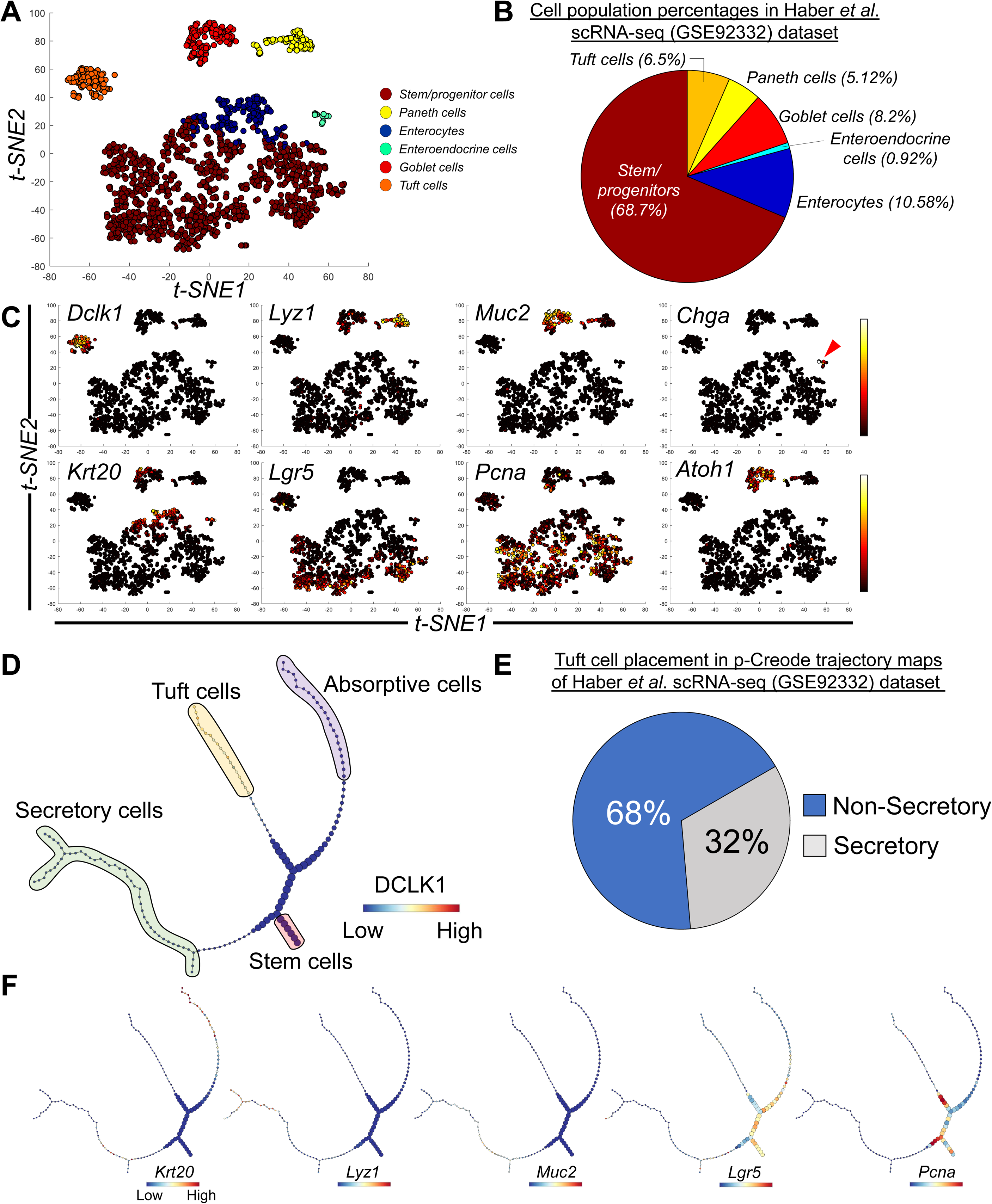
p-Creode analysis of GSE92332 scRNA-seq dataset, Related to Figure 2. **(A)** t-SNE analysis of the GSE92332 scRNA-seq dataset generated from the wildtype ileal epithelium. Cell type clusters, including goblet cells (red), Paneth cells (yellow), enteroendocrine cells (green), tuft cells (orange), enterocytes (dark blue), and stem/progenitor cells (brown), were identified by k-means clustering and manually annotated. Each datapoint represents a single cell from a 1,522 cell-dataset. **(B)** Quantification of cell populations within the GSE92332 datasets. Tuft cell (orange) percentage is 6.5% of the 1,522 cell-dataset. **(C)** Overlay of selected cell identity genes, including, including *Dclk1*, *Lyz1*, *Muc2*, *Chga*, *Krt20*, *Lgr5*, *Pcna*, and *Atoh1*, on the wildtype t-SNE plot. Red arrows in the *Chga* plot denote the enteroendocrine cluster. **(D)** Representative p-Creode map of GSE92332 scRNA-seq dataset. Graph overlay depicts *Dclk1*. Cell lineages, including secretory cells (green), absorptive cells (purple), tuft cells (orange), and stem cells (red), were manually labelled. Node size represents cell state density and each edge represents cell state transitions. Overlay represents ArcSinh-scaled gene expression data. **(E)** Quantification of n = 100 p-Creode maps show that 68% of graphs are classified as non-secretory and 32% as secretory. **(F)** Overlay of selected cell identity genes, including *Chga*, *Lyz1*, *Muc2*, *Krt20*, *Lgr5*, and *Pcna*, depicting epithelial cell differentiation in the wildtype topology. Overlays represent ArcSinh-scaled gene expression data.

**Figure S9.**
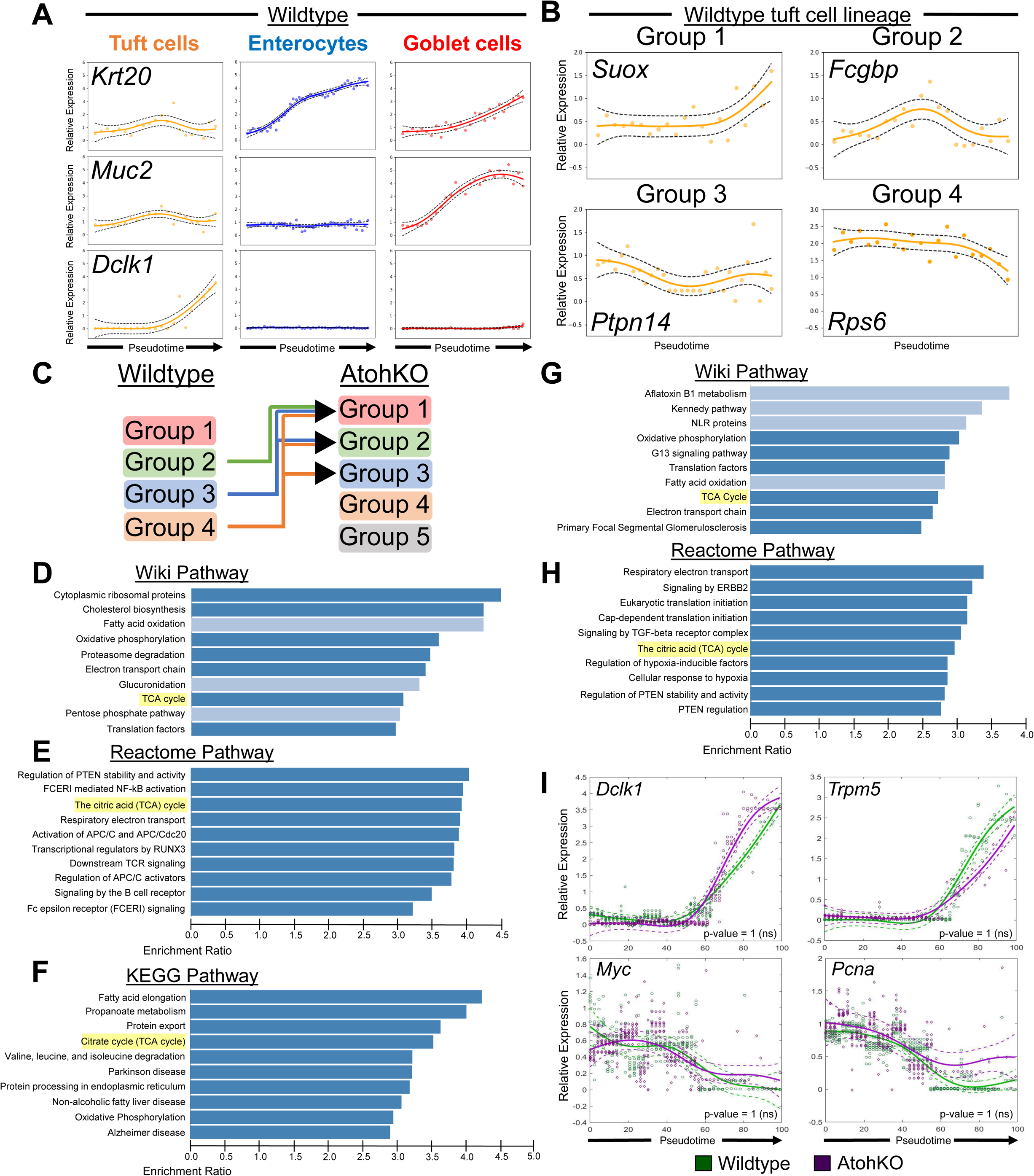
Trend dynamic analysis for p-Creode cell lineages, Related to Figure 3. **(A)** Relative expression of cell identity genes, *Krt20, Muc2,* and *Dclk1,* along the wildtype tuft cell (orange), enterocyte (blue), and goblet cell (red) lineages along pseudotime. Gene expression is represented by ArcSinh-scaled data. **(B)** Representative gene trends (solid orange line) across pseudotime from Groups 1 (e.g. *Suox*), 2 (e.g. *Fcgbp*), 3 (e.g. *Ptpn14*), and 4 (e.g. *Rps6*) of trend dynamic clustering of the tuft cell lineage. Solid circles depict raw data from single representative p-Creode graph and dashed black line represents the confidence interval for gene expression. **(C)** Schematic depicting the comparison between wildtype and AtohKO gene clusters. We identified 1,755 genes that class switched from wildtype groups 2-4 to AtohKO groups 1-3. **(D-E)** Over-representation analysis using **(D)** Wiki and **(E)** Reactome Pathway datasets of 5-trend analysis. **(F-H)** Over-representation analysis using **(F)** KEGG, **(G)** Wiki Pathway, and **(H)** Reactome Pathway datasets of 3-trend analysis. **(I)** Trend dynamics along pseudotime of non-class switching tuft cell-expressing genes, *Dclk1* and *Trpm5*, and stem cell-expressing genes, *Myc* and *Pcna*, between wildtype (green) and AtohKO (magenta). Solid line represents wildtype (green) and AtohKO (magenta) gene expression trends from 10 representative p-Creode graphs. Raw data is shown for wildtype (green circles) and AtohKO (magenta diamonds). Confidence interval of raw data is depicted by dashed lines for wildtype (green) and AtohKO (magenta). Dynamic time warping was used to fit wildtype and AtohKO tuft cell data to the same scale. Statistical analysis of trend differences and consensus alignment was performed between conditions, p-value = 1, not significant (ns) by t-test.

**Figure S10.**
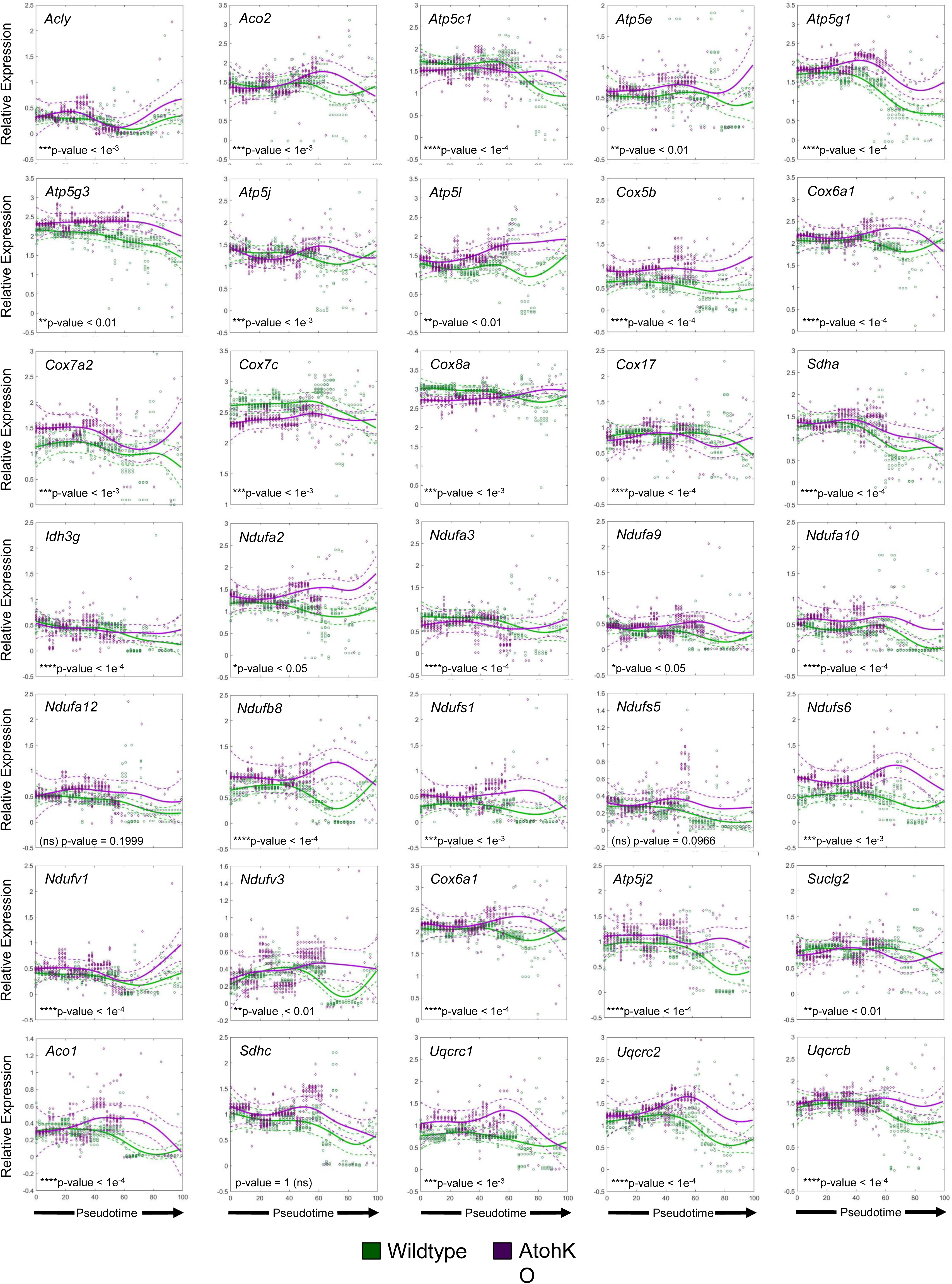
TCA cycle genes in the wildtype and AtohKO tuft cell lineage, Related to Figure 3. Trend dynamics along pseudotime of TCA cycle-associated genes between wildtype (green) and AtohKO (magenta). Solid line represents wildtype (green) and AtohKO (magenta) gene expression trends from 10 representative p-Creode graphs. Raw data is shown for wildtype (green circles) and AtohKO (magenta diamonds). Confidence interval of raw data is depicted by dashed lines for wildtype (green) and AtohKO (magenta). Dynamic time warping was used to fit wildtype and AtohKO tuft cell data to the same scale. Statistical analysis of trend differences and consensus alignment was performed between conditions, *p-value < 0.05, **p-value < 0.01, ***p-value < 0.001, ****p-value < 0.0001, and not significant (ns) by t-test.

**Figure S11.**
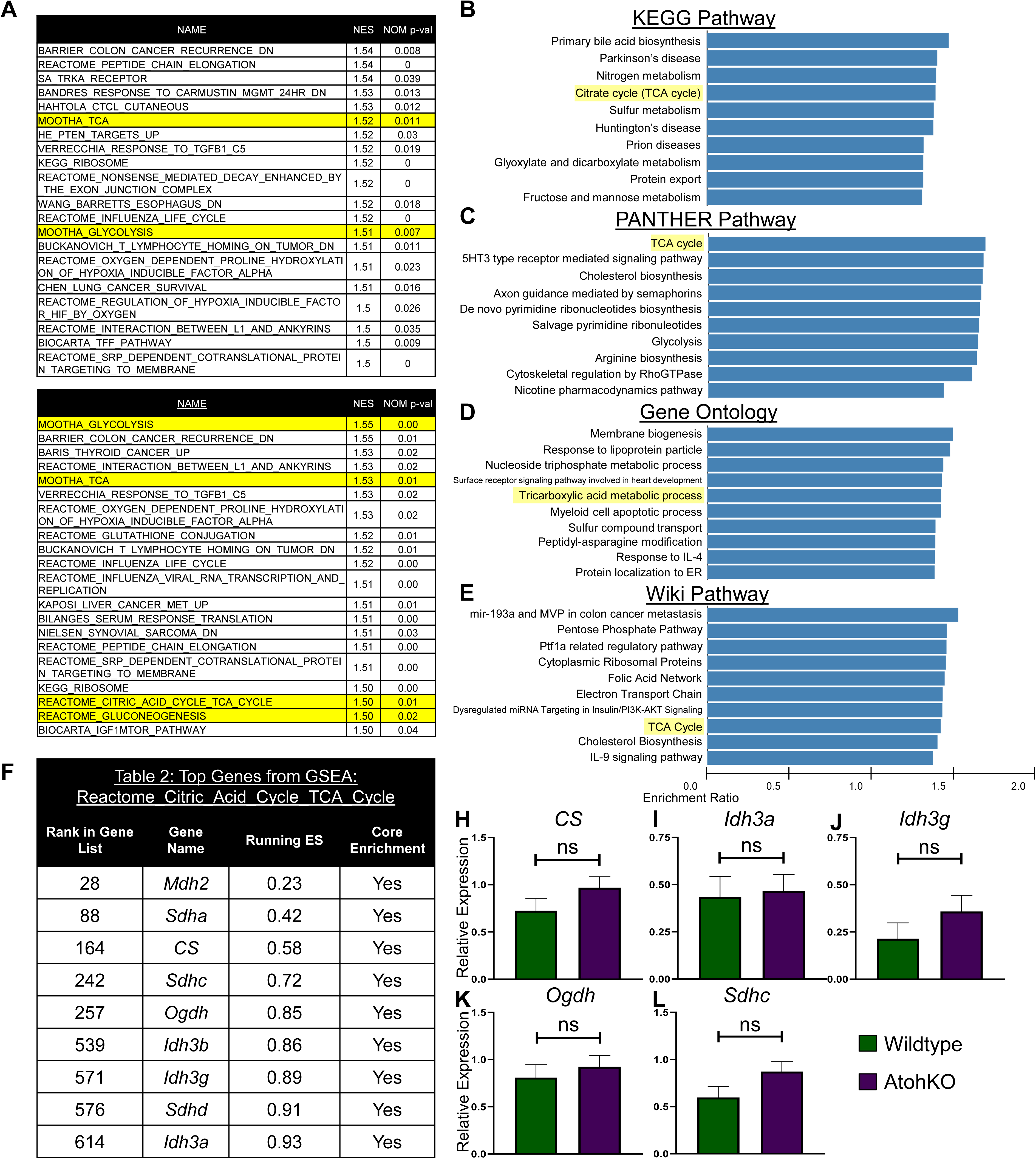
Gene expression analysis of wildtype and AtohKO tuft cells, Related to Figure 3. **(A)** Two independent GSEA iterations of median difference in gene expression between wildtype and AtohKO tuft cells. Top 20 gene sets from positive gene enrichment are ranked by the normalized enrichment score (NES) and p-value. Yellow highlighted gene sets are related to the citric acid cycle and metabolism pathways. **(B-E)** Functional analysis between wildtype and AtohKO tuft cells for **(B)** KEGG pathway, **(C)** PANTHER pathway, **(D)** Gene Ontology, and **(E)** Wiki Pathway analysis shows positive enrichment for the tricarboxylic acid cycle (yellow highlighted). **(F)** Ranked gene list from the metabolism-related gene list “Reactome Citric Acid Cycle TCA Cycle.” Genes contributing to the running enrichment score (ES) are labeled “Yes” for Core Enrichment. **(H-L)** Relative expression of TCA cycle genes in wildtype (green) and AtohKO (magenta) tuft cells. Error bars represent SEM from wildtype (n = 58 cells) and AtohKO (n = 64 cells). Not significant (ns) by t-test.

**Supplementary Figure 12.**
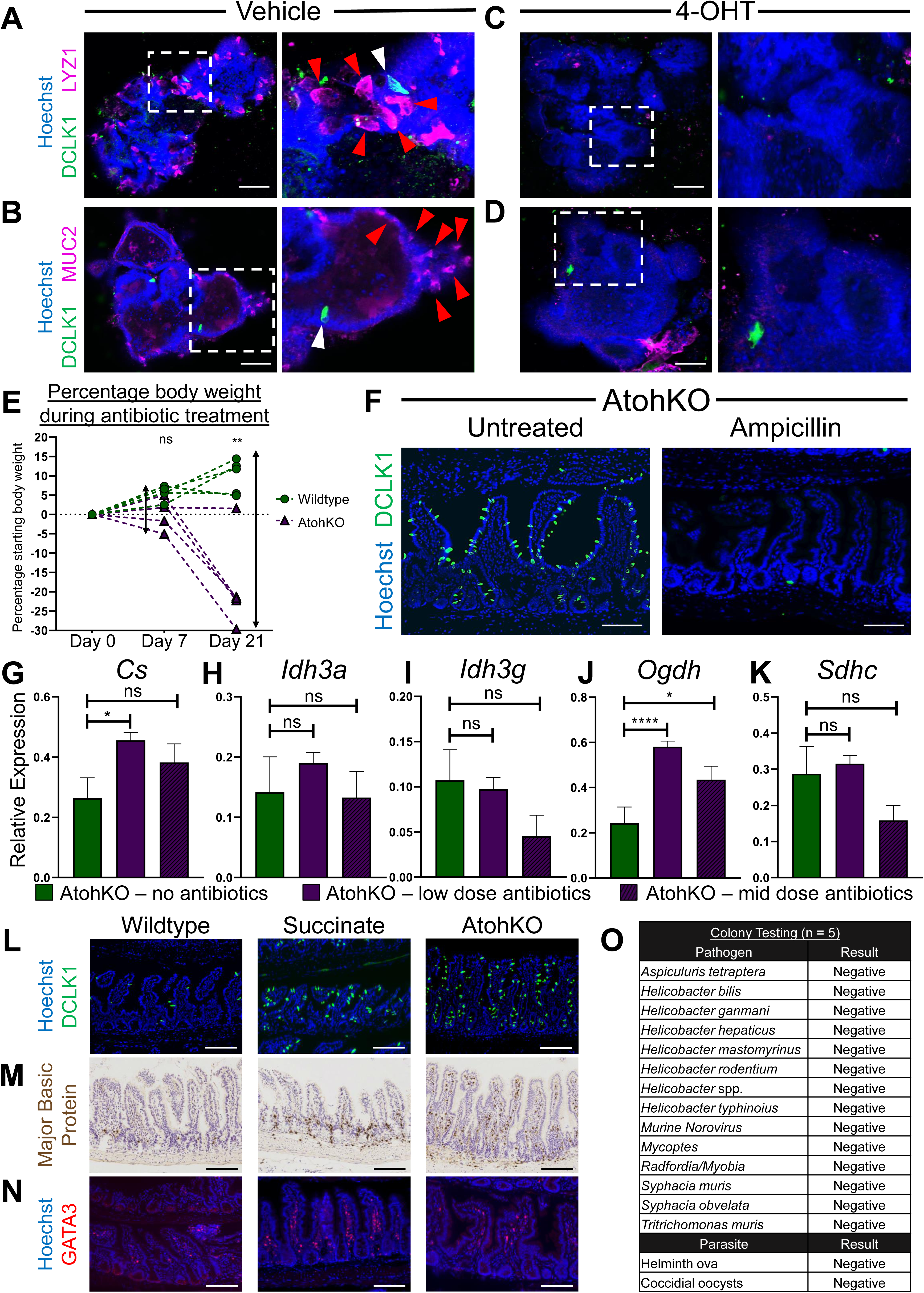
Microbiome contribution is necessary for AtohKO tuft cell hyperplasia, Related to Figure 4. **(A)** Vehicle-treated *Lrig1^CreERT2/+^;Atoh1^fl/fl^* small intestinal enteroids are positive for DCLK1+ tuft cells (green) and LYZ1+ Paneth cells (magenta). Magnified inset (white dashed box) shows LYZ1-positive Paneth cells (red arrows) and DCLK1+ tuft cell (white arrow). Hoechst (blue) denotes nuclei, scale bar = 50 µm. **(B)** Vehicle-treated *Lrig1^CreERT2/+^;Atoh1^fl/fl^* small intestinal enteroids are positive for DCLK1+ tuft cells (green) and MUC2+ goblet cells (magenta). Magnified inset (white dashed box) shows MUC2-positive Paneth cells (red arrows) and DCLK1+ tuft cell (white arrow). Hoechst (blue) denotes nuclei, scale bar = 50 µm. **(C-D)** 4-OHT-treated *Lrig1^CreERT2/+^;Atoh1^fl/fl^* small intestinal enteroids lack DCLK1+ and both **(C)** LYZ1+ Paneth (magenta) and **(D)** MUC2+ goblet cells (magenta). Hoechst (blue) denotes nuclei, scale bar = 50 µm. **(E)** Percentage change in body weight during antibiotic and tamoxifen treatment. Day 0 – start of antibiotic treatment, Day 7 – start of tamoxifen treatment (antibiotics continued), and Day 21 – cessation of tamoxifen (and antibiotic treatment). Wildtype (green) (n = 5 replicates) and AtohKO (magenta) (n = 5 replicates). **p < 0.01 and not significant (ns) by t-test. **(F)** Immunofluorescence imaging of DCLK1 (green) in untreated and Ampicillin-treated (1 g/ml) AtohKO small intestine. Hoechst (blue) denotes nuclei. Scale bar = 100 µm. **(G-K)** Relative expression of tricarboxylic acid cycle-related genes in AtohKO – no antibiotics (green), AtohKO – low dose antibiotics (magenta), and AtohKO – mid dose antibiotics (magenta, dashed) tuft cells. Error bars represent SEM from untreated AtohKO (n = 25 cells), low-dose antibiotic-treated AtohKO (n = 202 cells), and mid-dose antibiotic-treated AtohKO (n = 36 cells). ****p < 0.001, *p < 0.05, and not significant (ns) by t-test. **(L)** Immunofluorescence imaging of DCLK1 (green) in wildtype, succinate-treated, and AtohKO small intestine. Hoechst denotes nuclei. Scale bar = 100 µm. **(M)** Immunohistochemical staining of major basic protein (brown) in wildtype, succinate-treated, and AtohKO small intestine. Blue denotes nuclei. Scale bar = 100 µm. **(N)** Immunofluorescence imaging of GATA3 (red) in wildtype, succinate-treated, and AtohKO small intestine, Hoechst denotes nuclei. Scale bar = 100 µm. **(O)** Composite of colony testing (n = 5 mice) for pathogenic helminth and eukaryotic protists.

**Figure S13.**
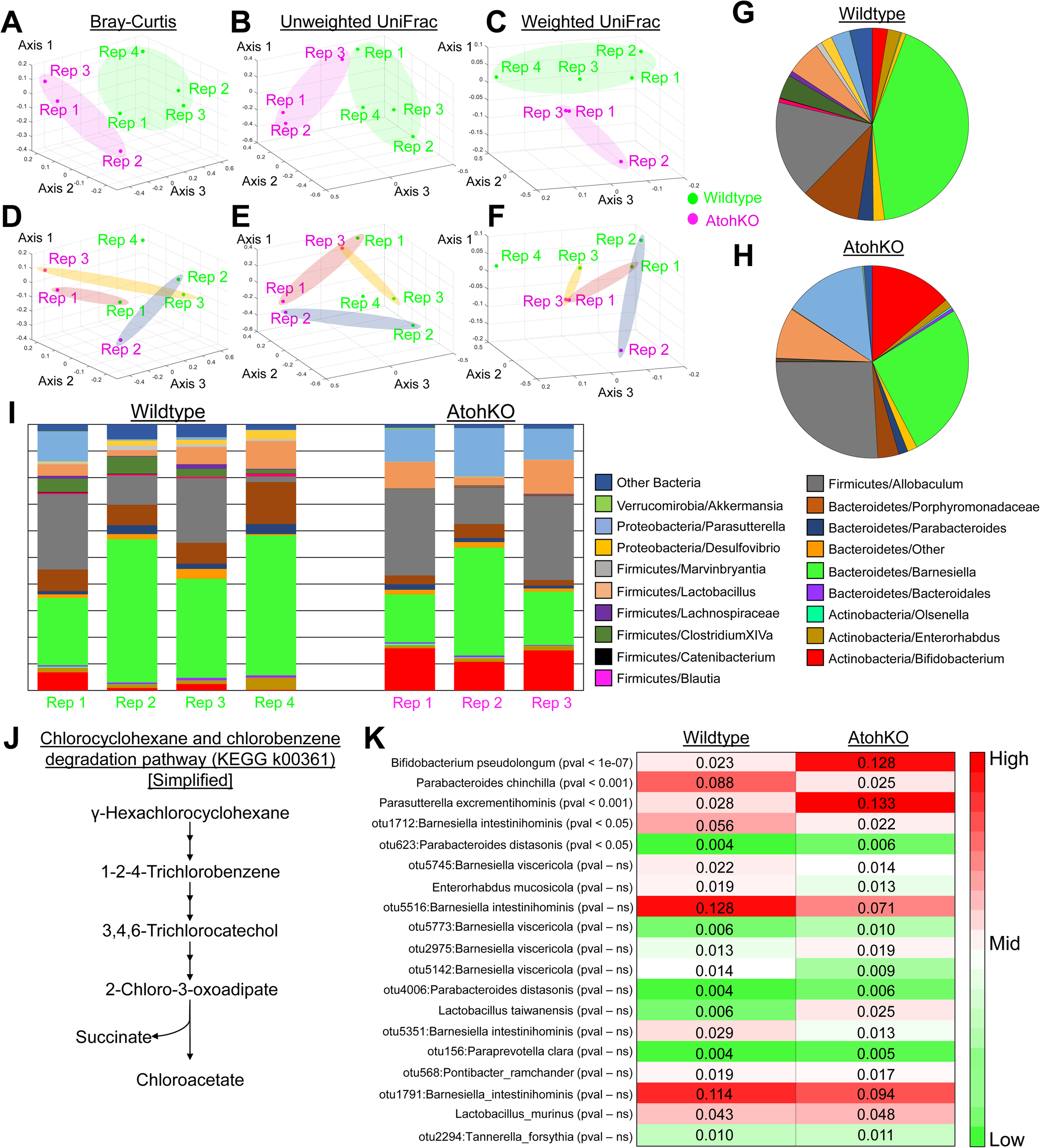
Functional analysis of microbiome profile reveals enrichment of succinate producing bacteria in the AtohKO ileum, Related to Figure 4. **(A-F)** Principal component analysis of ileal microbiome profiles as determined by **(A, D)** Bray-Curtis dissimilarity, **(B, E)** Unweighted UniFrac, and **(C, F)** Weighted UniFrac based on 16s rRNA amplicon sequencing. Annotation of PCA plots indicates clustering of wildtype (n = 4, green) and AtohKO (n = 3, magenta) biological replicates by **(A-C)** phenotype and **(D-F)** co-housing. **(G-H)** Average relative abundance in wildtype (n = 4) and AtohKO (n = 3) replicates at the genus level. **(I)** Relative abundance from each wildtype (green, n = 4) and AtohKO (magenta, n = 3) replicate. Annotation reflects bacterial taxonomy at the genus level. **(J)** Heatmap of relative abundance at the operational taxonomic unit (OTU) level between wildtype and AtohKO. P-value calculated by t-test.

**Figure S14.**
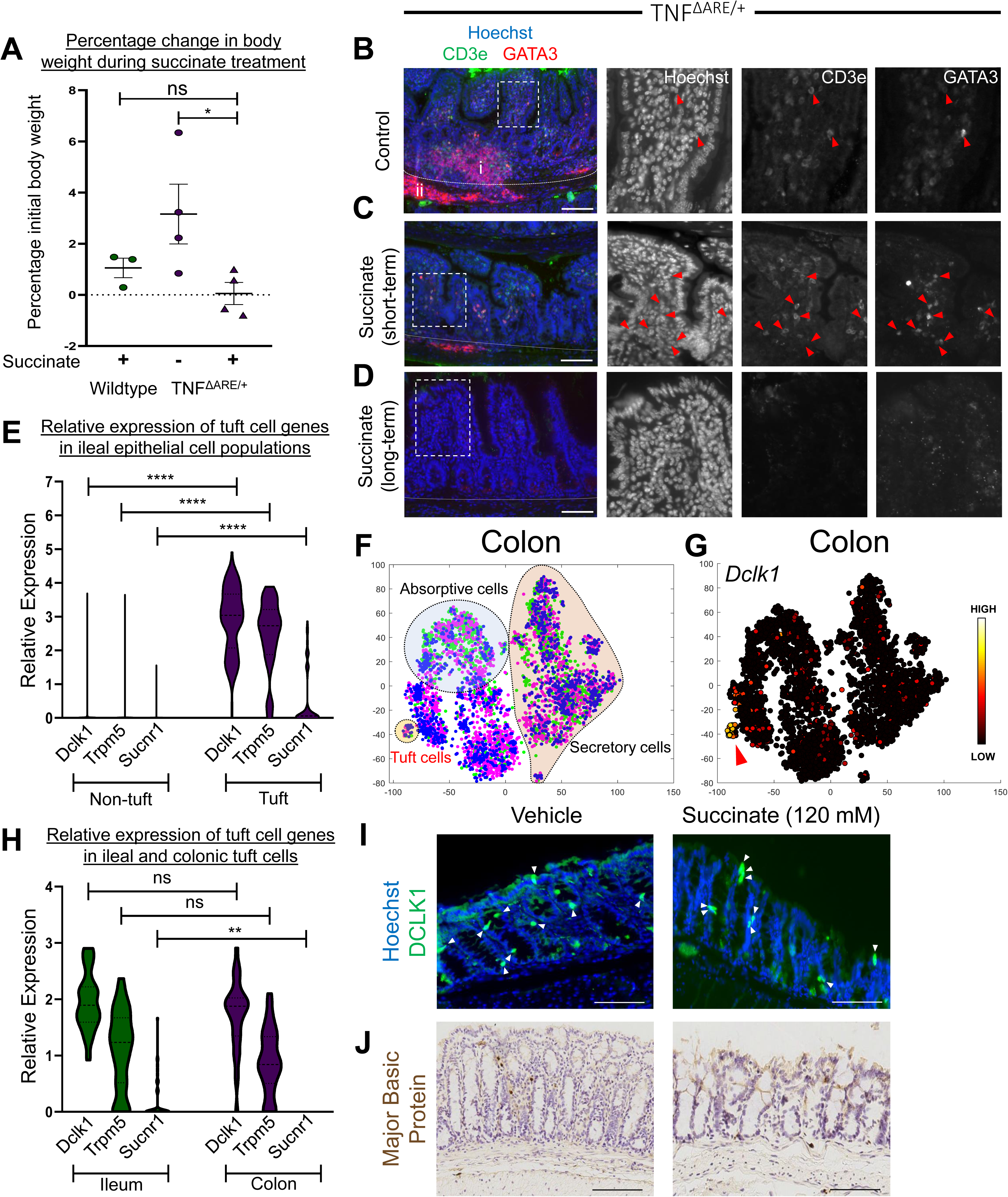
SUCNR1 is highly expressed in small intestinal tuft cells, Related to Figure 5. **(A)** Percentage change in body weight in untreated TNF^ΔARE/+^ (green circles) and succinate-treated wildtype (magenta circles) and TNF^ΔARE/+^ animals (magenta triangles). Error bars represent SEM across n = 4 wildtype replicates, n = 3 succinate-treated wildtype replicates, and n = 4 succinate treated-TNF^ΔARE/+^ replicates. *p < 0.05 and not significant (ns) by t-test. **(B-D)** Immunofluorescence imaging of CD3e (green) and GATA3 (red) in **(B)** untreated, **(C)** short-term, and **(D)** long-term succinate treated TNF^ΔARE/+^ animals. Hoechst (blue) denotes nuclei, scale bar = 100 µm. Single channel images are shown for magnified inset (white box) and dashed white line demarcates the stromal region. Red arrows denote CD3e+ GATA3+ lymphocytes. **(E)** Quantification of relative expression of *Dclk1*, *Trpm5,* and *Sucnr1* in non-tuft (green) and tuft (magenta) cells from the wildtype ileal epithelium. Expression data is generated from 6,862 non-tuft cells and 70 tuft cells across six wildtype replicates. **(F)** t-SNE analysis of wildtype colonic scRNA-seq dataset consisting of 3,980 cells. Manual annotation of the t-SNE plot demonstrates the absence of segregation in datapoints from three biological replicates (Rep. 1 – green, Rep. 2 – magenta, and Rep. 3 – blue). Each datapoint represents a single cell. The absorptive (blue), secretory (red), and tuft cell (orange) populations are demarcated. **(G)** *Dclk1* expression in wildtype colonic epithelial cells. Overlay represents ArcSinh-scaled expression data. **(H)** Quantification of relative expression of *Dclk1*, *Trpm5*, and *Sucnr1* in ileal (green) and colonic (magenta) tuft cells. Expression data is generated from 58 wildtype tuft cells (n = 4 ileal replicates) and 32 wildtype colonic tuft cells (n = 3 colonic replicates). **p < 0.01, not significant (ns) by t-test. **(I)** Immunofluorescence imaging of DCLK1 (green) vehicle and succinate-treated colon. Hoechst (blue) denotes nuclei and white arrows denote tuft cells. Scale bar = 100 µm. **(J)** Immunohistochemistry staining of major basic protein in vehicle and succinate-treated colon. Blue denotes nuclei and brown represents the counterstain. Scale bar = 100µm.

